# Nicotinamide reverses the Warburg effect in Chinese hamster ovary cell culture

**DOI:** 10.1101/2025.06.18.660349

**Authors:** James Morrissey, Ayca Cankorur-Cetinkaya, Luigi Grassi, Annie J. Harwood-Stamper, Jonathan Welsh, Cleo Kontoravdi

## Abstract

The Warburg effect, the preferential conversion of glucose-derived pyruvate to lactate despite available oxygen, is a key feature of Chinese hamster ovary (CHO) cell culture. Lactate accumulation in recombinant protein-producing cell culture is an inefficient usage of glucose, as well as being deleterious to cells. Lactate accumulation lowers culture pH, requiring base addition to maintain bioreactor pH setpoint, which subsequently leads to hyperosmolarity, adversely impacting cell growth, productivity and product quality. A key driver for the Warburg effect, and hence lactate accumulation, is the need to regenerate NAD^+^ consumed during glycolysis. Since oxidative phosphorylation (OXPHOS) has limited capacity to recycle NADH back to NAD^+^ at high glycolytic fluxes, cells rely on lactate dehydrogenase (LDH) to convert pyruvate to lactate, simultaneously regenerating NAD^+^ and sustaining glycolysis. Thus, providing the cells capacity to generate more NAD^+^ would decrease the reliance on the Warburg effect. In this study, feeding the NAD^+^ precursor nicotinamide (NAM) leads to reversal of the Warburg effect, inducing the “lactate shift” three days earlier in cell culture and reducing peak lactate concentration by 40%. Transcriptomic analysis further confirms this metabolic shift, with an upregulation of key mitochondrial electron transport chain genes. These results identify NAD+/NADH balance as a key regulator of the Warburg effect and demonstrate NAM supplementation as a simple, cost-effective strategy to mitigate lactate accumulation and improve metabolic efficiency in CHO cell cultures.

## 1 Introduction

Chinese hamster ovary (CHO) cells are the workhorse of biopharmaceutical production (Walsh and Walsh, 2022). CHO cells offer several advantages, including the capacity for post-translational modifications similar to those in human cells, which is crucial for the efficacy and safety of biopharmaceutical products. Over the years, CHO cells have been optimised for higher yield, stability, and scalability, making them the preferred choice for the production of biopharmaceuticals such as monoclonal antibodies (Kunert and Reinhart, 2016).

A key challenge still faced by the biopharmaceutical industry is the accumulation of lactate in cell culture. Lactate accumulation is a result of aerobic glycolysis, also known as the Warburg effect, which is the conversion of glucose to lactate despite the presence of oxygen. The Warburg effect is characteristic of CHO cells (Hartley et al., 2018), which typically secrete lactate during the exponential (growth) phase of cell culture, followed by a lactate switch where the accumulated lactate is then consumed by the cells. This lactate switch is a desired characteristic of cell culture and is linked to a healthy and more oxidative cell phenotype (Mulukutla et al., 2012), therefore encouraging a lactate switch can lead to higher titres and harvest viability.

Lactate secretion as a result of the Warburg effect is both inefficient and deleterious to cell cultures. The production of ATP through glycolysis only yields 2 ATP per glucose molecule, compared to the ∼32 ATP generated by oxidative phosphorylation (OXPHOS). Re-routing the glucose-derived pyruvate away from lactate production and into the TCA cycle and electron transport chain (ETC), would be a more effective utilisation of glucose and, in theory, could increase cell growth in the early stages of cell culture and boost final product titres.

As well as being an inefficient use of glucose, lactate itself has a negative impact on cell culture. When lactate levels rise to high concentrations, detrimental consequences on growth and productivity may arise (Buchsteiner et al., 2018; Fu et al., 2016; Hefzi et al., 2025; Lao and Toth, 1997; Li et al., 2012; Torres et al., 2018). Lactate also has an indirect deleterious impact on cell culture,through the drop in pH upon accumulation in the extracellular environment. In a pH-controlled bioreactor, lactate secretion lowers the pH below setpoint, causing the addition of base. This base addition will raise osmolality, and lead to inhibition of cell growth and productivity (Alhuthali et al., 2021; Romanova et al., 2022; Zhu et al., 2005)

Multiple explanations exist for CHO cells preference for aerobic glycolysis, yet the underlying mechanisms are yet to be fully elucidated. Amongst these explanations are a rapid need for ATP generation (Liberti and Locasale, 2016; Zheng, 2012), synthesis of biomass precursors (Boroughs and Deberardinis, 2015; Cairns et al., 2011; Heiden et al., 2009), impaired mitochondrial function (Heiden et al., 2009; Luengo et al., 2021; Sacco et al., 2023) and proteome constraints (Sánchez et al., 2017; Yeo et al., 2020). Following lactate accumulation in the exponential phase, multiple factors then influence the lactate switch, including bioreactor pH (Becker et al., 2019; Zalai et al., 2015), glucose and glutamine concentration (Ghorbaniaghdam et al., 2014; Wahrheit et al., 2014).

Another key driving force for aerobic glycolysis is the balance of redox co-factors NAD^+^ and NADH. Lactate production and secretion is controlled by a single reaction, lactate dehydrogenase (LDH), which reversibly converts pyruvate to lactate, simultaneously regenerating a molecule of NAD^+^ from NADH. In the exponential phase of cell culture, CHO cells consume large amounts of glucose leading to high glycolytic fluxes, which consumes large amounts of NAD^+^ in the GAPDH reaction (Lunt and Vander Heiden, 2011). In order to regenerate NAD^+^ to maintain glycolytic function, cells rely on LDH and hence lactate is secreted as a by-product.

Increasing intracellular NAD^+^ concentrations may offer an effective strategy for reducing lactate secretion or even promoting lactate consumption. One approach involves adding NAD^+^ directly to the growth medium (Lee et al., 2023); however, this method is prohibitively expensive and results in reduced cell growth, rendering it unsuitable for large-scale manufacturing. Alternative strategies include utilising NADH dehydrogenases (Diaz-Ruiz et al., 2018; Kim et al., 2014; Talla et al., 2020), NADH oxidases (Geueke et al., 2003; Titov et al., 2016; Vemuri et al., 2007), or inhibiting NAD^+^ consuming enzymes such as Poly(ADP-ribose) polymerases (PARPs) (Fang et al., 2014; Pirinen et al., 2014; Sarkar et al., 2023).

Another method to enhance NAD^+^ concentration is to target NAD^+^ biosynthesis pathways. There are three key pathways involved in NAD^+^ production: de novo biosynthesis from tryptophan, the Preiss–Handler pathway, and the salvage pathway (Denu, 2007). Enzymes within these pathways can be manipulated to boost NAD^+^ levels. For instance, nicotinamide phosphoribosyl transferase (NAMPT) catalyses the conversion of nicotinamide (NAM) to NAD^+^ via nicotinamide mononucleotide (NMN). Intracellular NAD^+^ levels can be increased through the use of small-molecule activators (Gardell et al., 2019; Yao et al., 2022) or NAMPT overexpression (Audrito et al., 2020; Wang et al., 2011). Since NAD^+^ acts as a feedback inhibitor of NAMPT (Burgos and Schramm, 2008; Takahashi et al., 2010), activating NAMPT helps drive the equilibrium towards NAD^+^ production, enhancing its intracellular concentration beyond baseline levels. Other enzyme targets in NAD^+^ biosynthesis, such as NMNAT1 (Fang et al., 2022; Liang et al., 2015; Rossi et al., 2018), NAPRT (Baldassarri et al., 2023), NRK1 (Cercillieux et al., 2022), and NADS (Hashida et al., 2016), which have not yet been tested in CHO cells at the time of writing.

A more practical and cost-effective strategy involves supplementing NAD^+^ biosynthesis precursors in CHO cell cultures. This has been tested in media formulations (Han et al., 2024), but not in feed design in fed-batch cultures to improve lactate profiles. In mammals, tryptophan, nicotinic acid (NA), and NAM are dietary sources, while extensive recycling of NAD^+^ from NAM in the salvage pathway maintains cellular NAD^+^ levels (Liu et al., 2018). Several potential precursors, such as NAM, NA, nicotinamide riboside (NR), nicotinate mononucleotide (NAMN), nicotinate adenine dinucleotide (NAAD), quinolinic acid (QA), and tryptophan, are available for supplementation. The utilisation of these pathways is tissue- and cell line-dependent (Liu et al., 2018), with prior transcriptomic data from the our CHO-K1-derived cell line reveals that the *de novo* biosynthesis pathway is inactive (*data not shown)* making tryptophan and QA ineffective as supplements. Among the remaining options, cost considerations at large-scale production favour NA and NAM, as they are inexpensive, unlike NR, NAMN, and NAAD, which are costly speciality chemicals and therefore not economically viable for large scale production. Consequently, NA and NAM were selected for this study.

## 2 Methodology

### 2.1 Cell-line and fed-batch conditions

A clonal cell line, transfected with a construct encoding glutamine synthetase (GS) and a human IgG1 monoclonal antibody, which was generated using the proprietary AstraZeneca CHO host cell line derived from CHO-K1, was used in this study.

Cells were seeded at density 0.9×10^6^ cells ml^-1^ in proprietary media at an initial working volume of 14mL in an Ambr®-15 system (Sartorius Stedim Biotech, Hertfordshire, UK). A two-part feed was added, with first feed occurring when cells reached a proprietary target density, and on even days going forward. Glucose was controlled at a defined range. Dissolved oxygen (DO) was controlled at 50%. The pH was controlled at 7.05±0.1. When the upper deadband was reached, CO_2_ was gassed, when the lower deadband was reached, base (sodium bicarbonate) was added. The temperature was controlled at 36.5 °C.

NAM and NA (Sigma-Aldrich, St. Louis, MO, USA) were added to the bioreactors from 1M stock solutions. An initial shake flask screening study (data not shown) was conducted to determine optimal feed concentrations and timings. Based on these findings, the final feeding strategy included 2mM and 5mM additions into each feed day, as well as a single 10mM addition on day 6. The stated NAM and NA concentrations represent the nominal bioreactor concentration immediately after feeding. The actual concentrations were not measured and may have exceeded these values if uptake was lower than the feeding rate. A combination of NAM & NA 2mM supplementation on each feed day was also implemented.

### 2.2 Measurements

Extracellular glucose and lactate concentrations were measured with a YSI 2900D Biochemistry Analyzer (YSI, Yellow Springs, Ohio, USA). Cell density and viability were measured by a Vi-Cell XR (Beckman Coulter, Indianapolis, Indiana, USA).

Quantification of amino acid concentrations in supernatant samples was completed using the AccuTag system on a Waters Acquity ultraperformance liquid chromatography (UPLC) system (Waters, Elstree, UK) according to manufacturer’s instructions. Supernatant samples were collected on harvest day for mAb titre quantification, using a protein A high-performance liquid chromatography (HPLC) on an Agilent 1260 Infinity series (Agilent Technologies, Santa Clara, CA, USA) by comparing peak size from each sample with a calibration curve.

Integral of viable cell density (IVCD), in 10^6^cells hr ml^-1^ was calculated according to Equation 1.

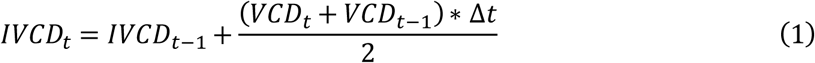

Where *VCD*_*t*_is the VCD at time t in 10^6^cells ml^-1^, Δ*t* is the time difference in hours.

Yield of lactate on glucose, *Y*_*lac*/*glc*_, was calculated using Equation 2:

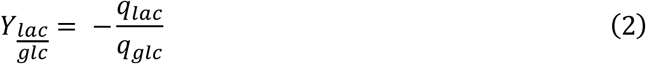

Cell specific productivity at time *t*, *qP*_*t*_, in pg cell^-1^ hr^-1^ was calculated using Equation 3:

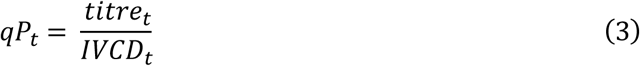

Statistical analysis on *Y*_*lac*/*glc*_ and *qP*_*t*_ to compare each feeding strategy with the control condition was performed using an unpaired two-tailed t-test in Microsoft Excel.

### 2.3 Transcriptomic Data

RNA was extracted from cell pellets containing 1×10^7^ cells, in accordance with the manufacture’s protocol of a RNeasy Mini Kit (Qiagen, Manchester, UK). 1×10^7^ cells were taken from bioreactors with varying experimental conditions on day 6 and stored using RNAlater (Thermo Fisher Scientific, Waltham, MA, USA). Total RNA was then extracted with the RNeasy Mini Kit according to manufacturer’s instructions. RNA quality was checked using a Bioanalyzer (Agilent, Santa Clara, CA, USA). Library preparation was conducted utilizing a TruSeq Stranded Total RNA Sample Preparation (Illumina, San Diego, CA, USA). Sequencing was performed at the Next Generation Sequencing Facility at the University of Leeds on a NextSeq 2000 sequencer (Illumina, San Diego, CA, USA).

Fastp (Chen et al., 2018), version 0.23.4, was used to trim PCR and sequencing adapters and filter low quality read pairs. Trimmed and qc filtered reads were then used by Salmon (Patro et al., 2017), version 1.10.0, with the selective alignment strategy (Srivastava et al., 2020) to quantify the expression of transcripts, annotated in Ensembl, version 93 (Howe et al., 2021), with the addition of the extra transgenes. The Bioconductor package txtimport (Soneson et al., 2016) has been used to summarise transcript expression in terms of gene expression.

Differential gene expression (DGE) analysis was conducted using DESeq2 (Anders and Huber, 2010) in R version 4.4.2. (Genes with expression less than 1 transcript per million (TPM) in each sample were filtered from the results). A gene was considered differentially expressed if the adjusted p value (p_adj_) was less than 0.01 and absolute value of the Log2 fold change (Log2FC) was above 0.3. As RNA-Seq data was taken on day 6, before the single-dose feeds were added, there were 9 control experiments, 6 experiments with NAM feeding (2mM and 5mM feeds) and 6 with NA feeding (2mM and 5mM feeds) for the purposes of DGE analysis. DESeq2 analysis separating the 2mM and 5mM NAM feeds revealed similar differential expression between these sets and control (Supplementary Figure S4), meaning they were combined together for DGE.

Functional gene enrichment was carried out using Gene Set Enrichment Analysis (GSEA) (Subramanian et al., 2007) using the GSEApy package (Fang et al., 2023) in Python. GSEA was performed using the WikiPathways 2024 Human database (Agrawal et al., 2024), with a pre-ranked approach. A minimum gene set size of 3 and a maximum of 100 were used, with 1000 permutations. Pathways with a False Discovery Rate (FDR) q-value below 0.05 were considered significant.

Principal component analysis (PCA) was performed using scikit-learn (Pedregosa et al., 2011) in Python. Log₂(TPM+1) expression values were standardised with StandardScaler, and PCA was conducted with “n_components=2”. The first two components were used to visualise variance between samples.

## 3 Results and discussion

### 3.1 Nicotinamide feed supplementation reverses the Warburg effect

Three NAM feed supplementation strategies were tested in triplicate using an Ambr-15 bioreactor system. As outlined in Table 1, these included two conditions with NAM feeding on each feed day at different concentrations (2mM and 10mM bioreactor concentration), and a single-dose NAM feed on day 6 at 10mM bioreactor concentration. Figure 1 illustrates the effects of these NAM feeding strategies on extracellular lactate and glucose concentrations, as well as the lactate yield on glucose.

**Figure 1:**
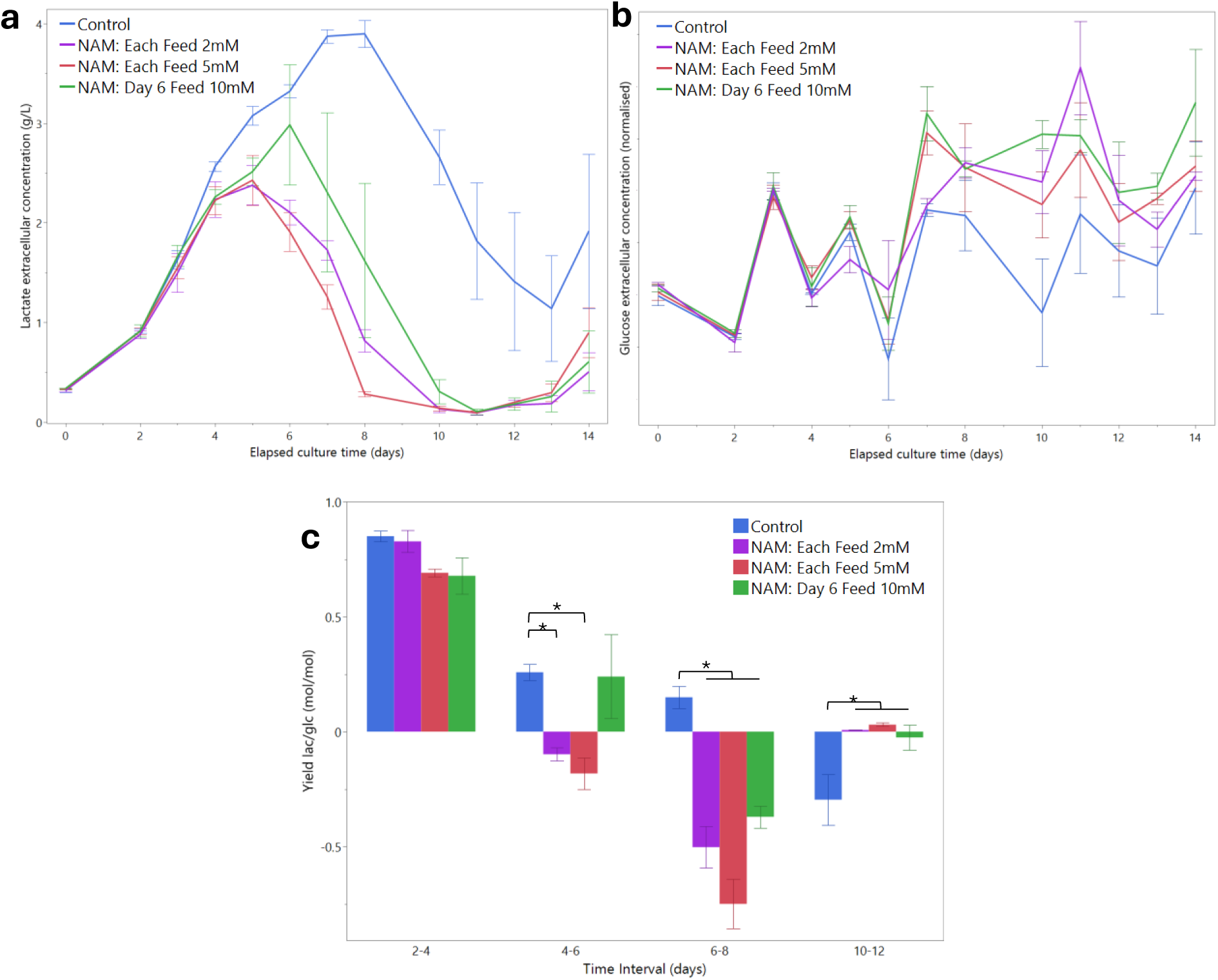
a) Lactate concentration, b) glucose concentration and c) yield of lactate on glucose (*Y*_*lac*/*glc*_) for the NAM-fed bioreactors. Error bars indicate standard error for the Ambr-15 vessel triplicates (n=3). Asterisk indicates statistical difference from control ( t-test p value <0.05). Glucose concentrations were normalised.

**Table 1:**
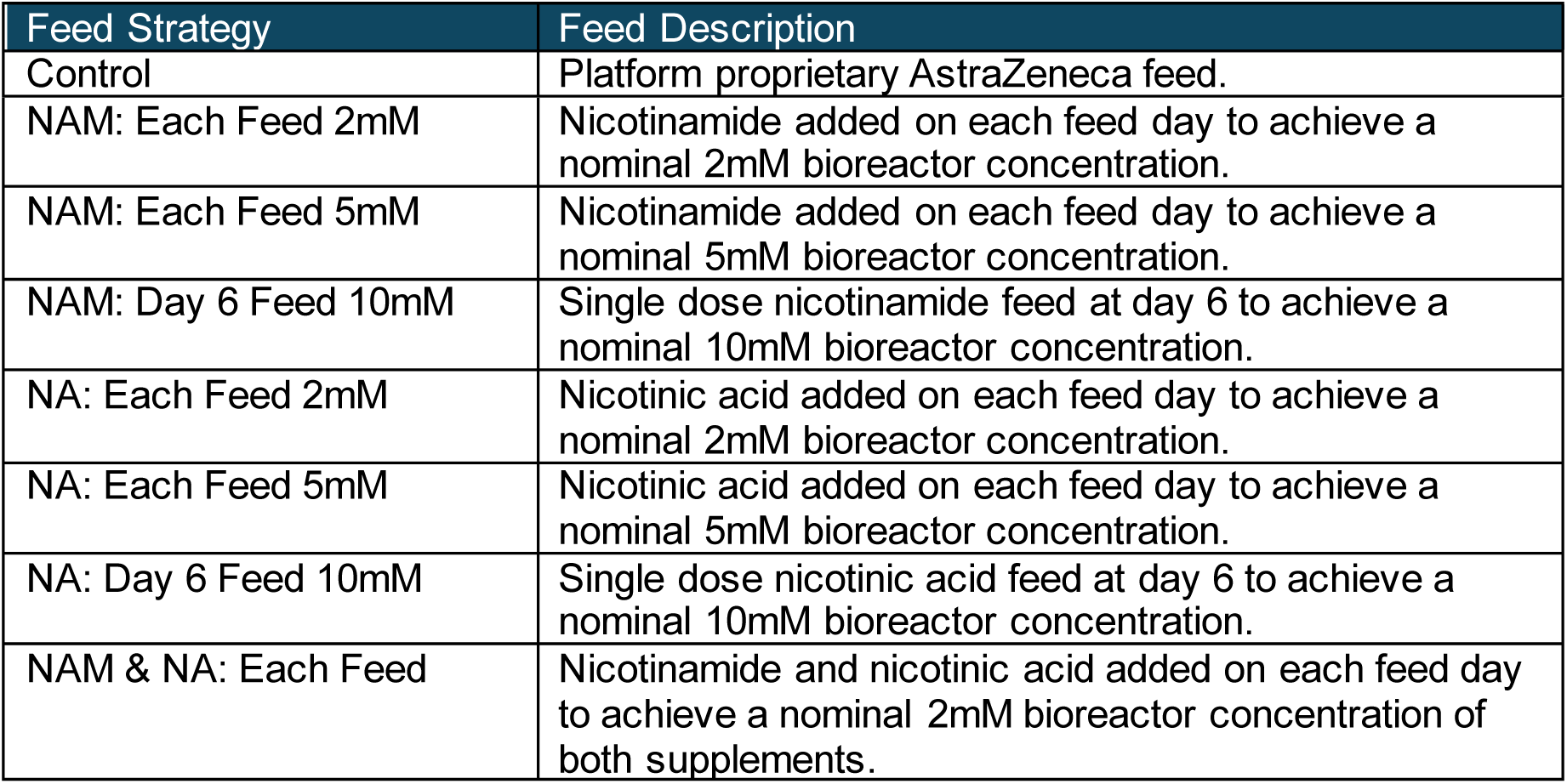
Summary of feeding strategies in Ambr-15. Each strategy was tested in triplicate.

| Feed Strategy | Feed Description |
| --- | --- |
| Control | Platform proprietary AstraZeneca feed. |
| NAM: Each Feed 2mM | Nicotinamide added on each feed day to achieve a nominal 2mM bioreactor concentration. |
| NAM: Each Feed 5mM | Nicotinamide added on each feed day to achieve a nominal 5mM bioreactor concentration. |
| NAM: Day 6 Feed 10mM | Single dose nicotinamide feed at day 6 to achieve a nominal 10mM bioreactor concentration. |
| NA: Each Feed 2mM | Nicotinic acid added on each feed day to achieve a nominal 2mM bioreactor concentration. |
| NA: Each Feed 5mM | Nicotinic acid added on each feed day to achieve a nominal 5mM bioreactor concentration. |
| NA: Day 6 Feed 10mM | Single dose nicotinic acid feed at day 6 to achieve a nominal 10mM bioreactor concentration. |
| NAM & NA: Each Feed | Nicotinamide and nicotinic acid added on each feed day to achieve a nominal 2mM bioreactor concentration of both supplements. |

Figure 1a demonstrates that NAM feed supplementation significantly reduced the average peak lactate concentration, lowering it from an average of 4.0 g L^-1^ in the control triplicates to 2.4 g L^-1^ in the 2mM and 5mM NAM-fed cultures (*p*=1.5×10^-4^), a 40% decrease in peak lactate concentration. As well as decreasing peak lactate, NAM feeding induced a shift to lactate consumption, with the lactate shift occurring on day 5 for both the 2mM and 5mM fed cultures, compared to day 8 in the control cultures.

Lactate concentration declined rapidly following this metabolic switch, indicating high rates of lactate consumption under these conditions. This effect is most pronounced for the single dose 10mM NAM feed, where lactate levels drop sharply after day 6. Figure 1c further demonstrates this, showing how *Y*_*lac*/*glc*_ turned negative (indicating lactate consumption) after NAM supplementation, while remaining positive in the control condition.

This early switch to lactate consumption is attributed to the conversion of NAM to NAD⁺ via the salvage pathway, mediated by NAMPT through NMN, leading to increased NAD⁺ concentrations (Han et al., 2024). The resulting elevated NAD⁺/NADH ratio strongly favours NAD^+^ consumption, converting lactate to pyruvate via LDH while generating NADH. Transcriptomic analysis on day 6 of cell culture, discussed further in Section 3.4, supports this metabolic shift, revealing significant upregulation of mitochondrial ETC genes (*Nd1*, *Nd2*, *Nd3*, *Nd4*, *Nd4l*, *Nd5*, *Cytb*, and *Apt6*), which suggests a transition towards OXPHOS. Additionally, the downregulation of *Hk2* indicates reduced glycolytic flux, reinforcing the observed decrease in glucose uptake. These transcriptomic changes align with the metabolic shift away from the Warburg effect.

In NAM-fed conditions, lactate serves as an alternative carbon source to glucose. As shown in Figure 1b, glucose uptake is reduced in the NAM-fed vessels, with lactate instead being utilised as a carbon source. This reduced glucose uptake leads to a consistent net flow of carbon toward pyruvate across all conditions, reflecting the highly regulated pathways downstream of pyruvate, such as the TCA cycle (Chong et al., 2010).

An additional factor contributing to the shift from glucose to lactate as a carbon source may be the regulatory impact of altered NAD⁺ metabolism. NAD⁺ is a crucial substrate for regulatory proteins like SIRTs and PARPs (D’Amours et al., 1999; Imai et al., 2000), and the increased NAD⁺ concentrations in NAM-fed conditions could impact glycolysis. For instance, PARP1 is known to inhibit hexokinase 1 (Fouquerel et al., 2014), suggesting that the heightened activity of these regulatory proteins may restrict glucose uptake. Additionally, NAM itself can inhibit SIRT activity (Bitterman et al., 2002; North and Verdin, 2004). SIRTs break down NAD⁺ into NAM during the deacetylation process and are auto-inhibited by NAM. This inhibition is especially significant for SIRT1 (Bitterman et al., 2002), which plays a key role in regulating glycolysis (Pinho et al., 2016; Rodgers et al., 2005). Therefore, the intracellular accumulation of NAM might block SIRT1 activity, potentially downregulating glycolysis and leading to reduced glucose uptake. SIRTs 1-3 activity also inhibits hypoxia-inducing factor-1α (*Hif1a*) (Greer et al., 2012), which regulates cellular response to hypoxia, by promoting glycolysis-related gene expression and suppressing OXPHOS. Further discussion and analysis of gene expression is found in Section 3.4.

### 3.2 High nicotinamide concentrations attenuate cell growth

Supplementation with NAM positively affects lactate profiles, but at high concentrations, it suppresses cell growth. As illustrated in Figure 2a, a NAM feed concentration of 5mM leads to a plateau in VCD starting from day 6 of culture and causes a more rapid decline in cell viability. The IVCD at harvest (day 14) for the control conditions averages at 5.3×10^12^cell hr ml^-1^, while it is 4.2×10^12^cell hr ml^-1^ for the 5mM NAM feed, a 20% decrease. These effects are less pronounced under conditions of lower NAM feed concentrations, (harvest IVCD at 2mM and single dose feed is 4.9 and 5.2×10^12^cell hr ml^-1^ respectively) suggesting a trade-off is needed between promoting a more oxidative state and affecting cell growth.

**Figure 2:**
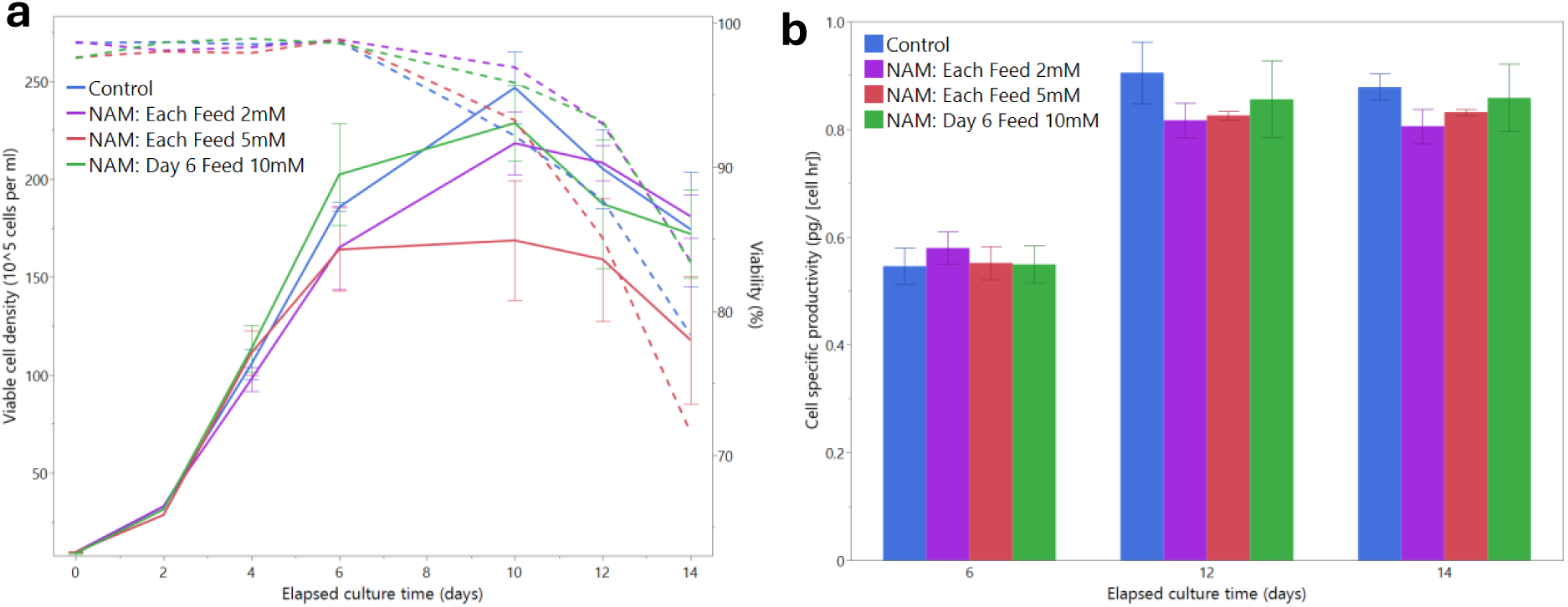
a) Viable cell density (VCD), viability (dashed line) and b) cell specific productivity for the recombinant production for the NAM-fed conditions. Error bars indicate standard error for the triplicates (n=3).

Several mechanisms may explain how NAM supplementation influences growth rates, and the lowered growth rate is supported by a downregulation of a host of growth-related genes which are discussed further in Section 3.4. One explanation is reduced glycolytic flux, resulting from the switch to lactate as a carbon source instead of glucose in the NAM-fed conditions. This glycolytic suppression can decrease cell proliferation, as glycolytic intermediates are vital precursors for nucleotide, amino acid, and lipid synthesis (Horváthová et al., 2021; Jones and Bianchi, 2015). With less glycolysis, fewer intermediates are available, compromising the synthesis of macromolecules essential for cell growth and division. For instance, glycolysis provides precursors like 3-phosphoglycerate, which is converted to serine and subsequently to glycine and cysteine, essential amino acids (Amelio et al., 2014). However, it has been shown that the high glycolytic fluxes are not essential to obtain biomass precursors, and a wild-type growth rate can be achieved in Warburg-null CHO cell lines (Hefzi et al., 2025).

Reduced glycolytic flux may contribute to an energy deficit, as the conversion of lactate to pyruvate via LDH does not generate ATP, unlike glycolysis, which produces two ATP molecules per glucose. However, the anticipated increase in OXPHOS capacity (see Section 3.4) may help offset this energy shortfall by enhancing ATP production through more efficient mitochondrial respiration. Furthermore, the synthesis of NAD^+^ from NAM through the salvage pathway imposes an additional ATP burden, requiring two ATP molecules per NAD^+^ molecule synthesised (Xie et al., 2020). Han et al. (2024) demonstrated that cells treated with NAD^+^ precursors exhibited lower ATP levels than untreated cells, reinforcing the idea that NAD^+^ biosynthesis increases cellular energy demand. Beyond ATP consumption, NAD^+^ synthesis from NAM consumes phosphoribosyl pyrophosphate (PRPP) as a key precursor, increasing the demand for nucleotides. The combined strain on ATP and nucleotide pools could explain the observed reduction in cell growth under high NAM concentrations.

Elevated NAD^+^ from NAM feeding also influences the NAD^+^ dependent regulatory proteins, such as SIRTs and PARPs, as discussed in Section 3.1, which may have negative effects on cell growth. Excessive PARP activity has been linked to DNA damage, genomic instability, and impaired cell proliferation (Kang et al., 2022). SIRTs, crucial for metabolic regulation, can be affected both by increased NAD^+^ and NAM itself. While NAD^+^ serves as a substrate for SIRT activation, NAM acts as a feedback inhibitor (Bitterman et al., 2002; North and Verdin, 2004). The concurrent rise in both activator and inhibitor levels could create unpredictable metabolic effects. Overactivity of SIRTs like SIRT1 and SIRT6, which deacetylate histones, could result in significant epigenetic changes that suppress genes required for growth and division. For instance, SIRT1 has been shown to inhibit the mTOR pathway, a key regulator of cell growth (Ghosh et al., 2010). Overactive mitochondrial SIRTs (SIRT3-5) may increase oxidative stress by enhancing mitochondrial metabolism, while NAM-induced inhibition of SIRTs may disrupt metabolic regulation, reducing ATP production and the availability of biosynthetic precursors necessary for growth. Additionally, SIRT1’s interaction with AMPK, a vital energy sensor, may be compromised with lowered SIRT1 activity, leading to metabolic dysregulation (Price et al., 2012).

Although high NAM concentrations inhibit growth, cell-specific productivity is not significantly affected (Figure 2b), indicating that productivity is not as reduced as drastically as growth rates. Since the impact of NAM on growth is dose-dependent, it is crucial to find a balance between its benefits and drawbacks. Implementing NAM supplementation in a platform process should involve optimising concentrations to avoid adverse effects on growth or titre (Supplementary Figure S1).

### 3.3 Nicotinic acid supplementation leads to lactate secretion spiral

NA feed supplementation did not yield the same beneficial outcomes as NAM. As shown in Figure 3a, NA-fed cultures initially maintained lactate levels comparable to the control but experienced a sharp increase in lactate concentration in the final days of culture. This lactate spike coincided with a decline in VCD and cell viability (Figure 3b). The excessive lactate secretion led to a decrease in pH, necessitating the addition of large volumes of base toward the end of the culture period. This base addition, in turn, triggered further lactate secretion, creating a ‘lactate secretion spiral’ observed in the NA-fed conditions.

**Figure 3:**
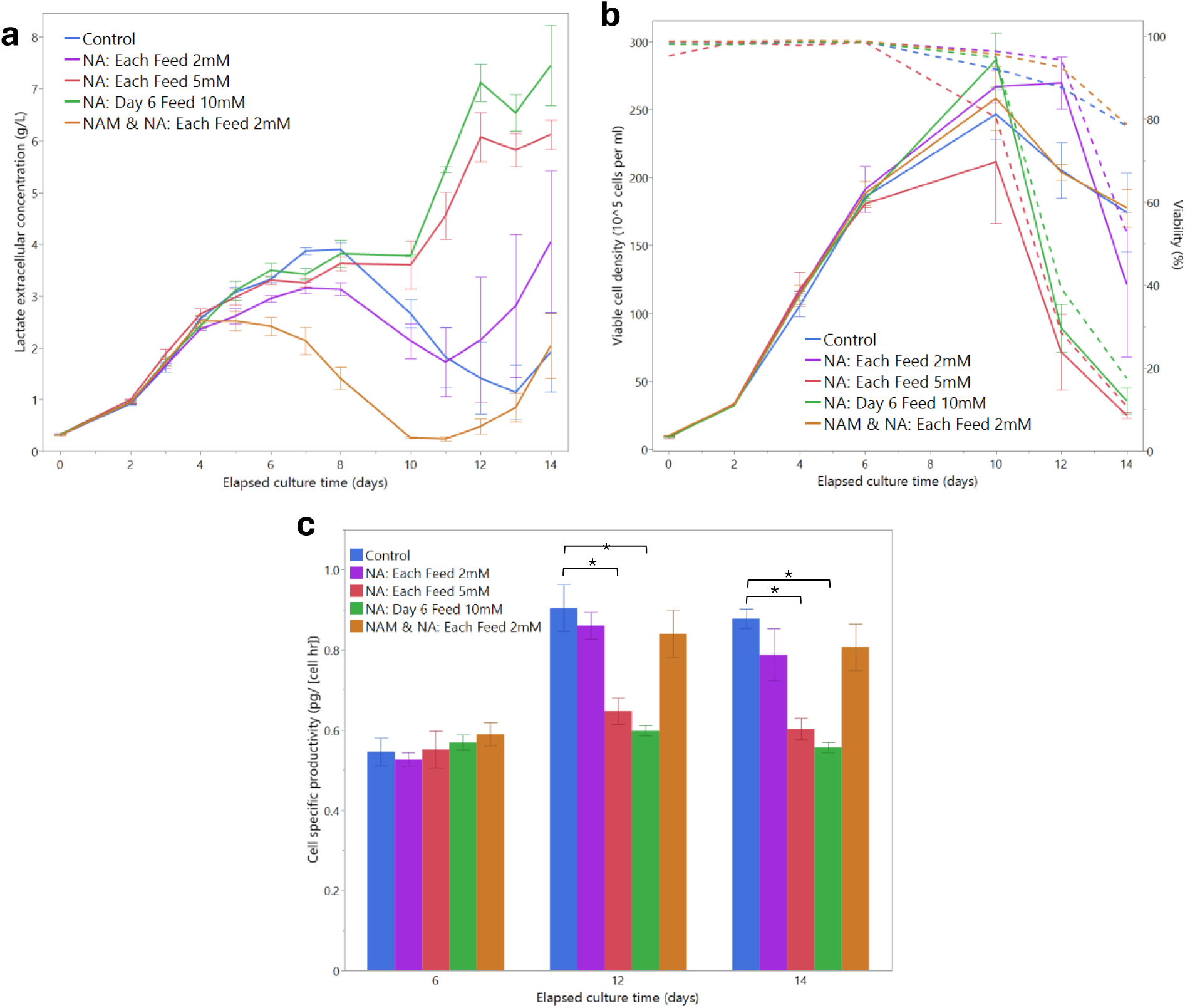
A) Lactate concentration, B) viable cell density (VCD), viability (dashed line) and C) cell specific productivity (qP) for the NA fed-conditions. Error bars indicate standard error for the triplicates. Asterisk indicates statistical difference from control (P value <0.05).

Even though culture vessels were pH-controlled, and the NA stock solution was buffered to neutral pH to minimise any direct impact on culture pH, the uptake of NA into cells may have caused a significant drop in intracellular pH beyond the capacity of acid-base transporters to regulate (Doyen et al., 2022). This intracellular acidification disrupts the mitochondrial proton gradient (Poburko et al., 2011), inhibits critical enzymes (Jubrias et al., 2003), and alters proton-dependent transport across the mitochondrial membrane (Selivanov et al., 2008).

The resulting lactate secretion sets off a secretion spiral in the following way: base is added to maintain the pH set point, which increases the osmolality beyond tolerable levels.

Elevated osmolality further impairs OXPHOS (Lee et al., 2003; Pan et al., 2019; Romanova et al., 2022), prompting even more lactate secretion and further base addition.

An additional complicating factor is the cellular uptake mechanism of NA. NA is transported into cells via the sodium-dependent MCT (SMCT) located on the plasma membrane (Bongarzone et al., 2020; Gopal et al., 2007, 2005; Yanase et al., 2008), which co-transports NA with one sodium ion. Since the base used in the culture is sodium bicarbonate, the increasing sodium concentration gradient from base addition enhances NA uptake via the SMCT. This further uptake is expected to exacerbate intracellular acidification, leading to more lactate secretion through MCT to counteract the pH imbalance. However, this response may be insufficient to maintain homeostasis, resulting in the observed decrease in cell viability.

### 3.4 Differential gene expression analysis confirms switch to oxidative metabolism

Transcriptomic analysis reveals a shift from glycolytic to oxidative metabolism, that provides further explanation to the reverse in the Warburg effect in NAM-fed cultures. Day 6 represents a crucial point in culture, as the control conditions were still secreting lactate, while the NAM-fed cells had just undergone the lactate switch. DGE analysis between control and NAM-fed using DESeq2 (Section 2.3) revealed 113 upregulated genes and 186 downregulated genes. Figure 4 displays a volcano plot of this DGE analysis, while Figure 5 (below) summarises the key impacts of NAM feeding on CHO cell metabolism. The full list of up and downregulated genes is available in Supplementary Table S1.

**Figure 4:**
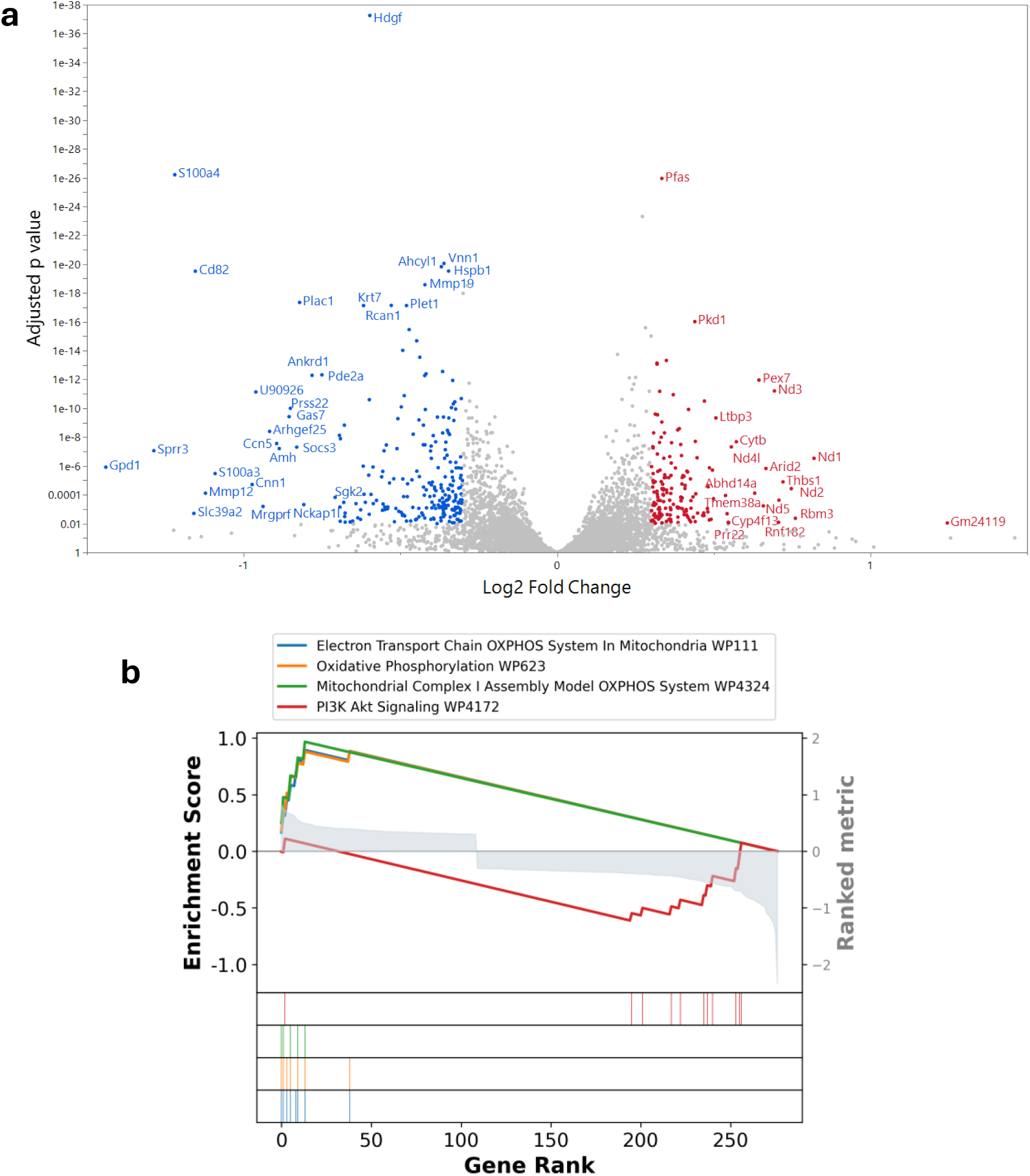
a) Volcano plot from differential gene expression (DGE) analysis between control (n=9) and NAM-fed (n=6) conditions at day 6. Upregulated genes (113) are shown in red, downregulated genes ( 186) in blue. Cutoff for differential expression is p_adj_ < 0.01 and abs(Log2 FC) > 0.3. b) GSEA plot showing enriched pathways from the DGE analysis.

**Figure 5:**
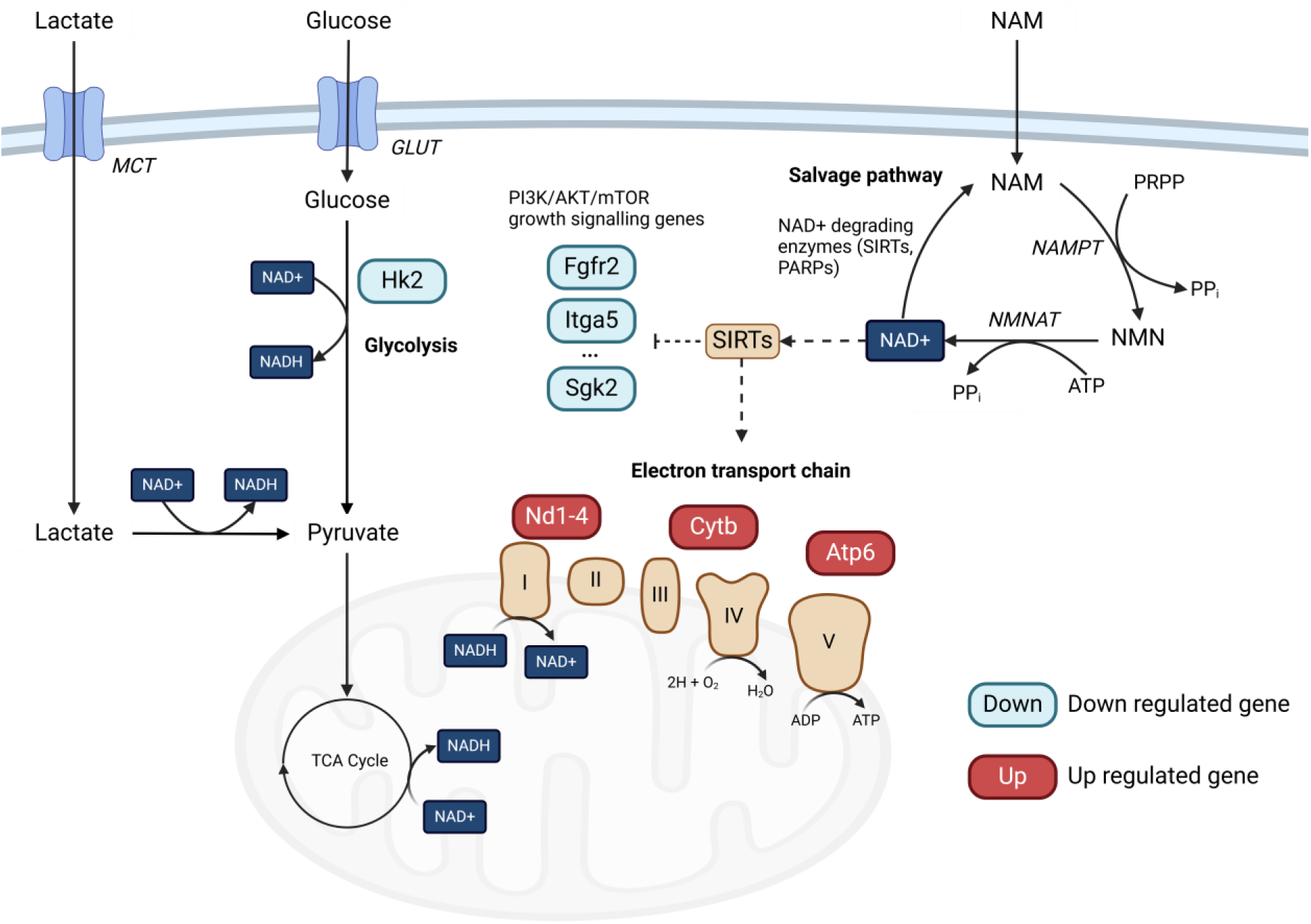
Summary of impact of NAM feeding. NAM is transported into the cell, converted to NMN and finally NAD+ in the salvage pathway. NAD+ impacts the Sirtuin class of regulatory proteins, impacting energy metabolism, including the electron transport chain and growth signalling genes which leads to a lower growth rate. As a result of increased NAD+, lactate is consumed alongside glucose, regenerating NADH in the LDH reaction. Glucose uptake and glycolysis is downregulated, while mitochondrial ETC genes are upregulated as cells rely on mitochondrial respiration for ATP. Red genes are upregulated, blue are downregulated, dashed lines indicate activation and blocked lines inhibition.

A key observation is the upregulation of multiple genes in the mitochondrial ETC, particularly *Nd1*, *Nd2*, *Nd3*, *Nd4*, *Nd4l*, *Nd5*, *Cytb*, and *Apt6*, which encode subunits of mitochondrial Complexes I, III, and V. This strongly suggests an increase in OXPHOS activity in NAM-fed cultures, consistent with the observed reduction in lactate secretion and the shift away from aerobic glycolysis.

One of the downregulated genes with the largest fold change is *Gpd1*(glycerol-3-phosphate dehydrogenase). *Gpd1* is part of the glycerol-3-phosphate shuttle, transferring NADH electrons to mitochondria and replenishing cytosolic NAD+. But with elevated NAD⁺ levels and reduced glycolysis, there is less NADH to be transported into the mitochondria and this gene is significantly downregulated.

The high expression of *Pfas* (phosphoribosylformylglycinamidine synthase), in NAM-fed conditions is likely driven by the increased demand for nucleotides in NAM-fed condition. Since NAM is converted to NAD⁺ via the salvage pathway, this process consumes both ATP and PRPP, a key substrate in de novo purine biosynthesis. To compensate, cells upregulate *Pfas*, a crucial enzyme in purine metabolism, to meet purine demand. The increased flux through this reaction may also explain the high glutamine consumption rates and accumulation of glutamate in NAM-fed conditions (Supplementary Figure S2), as *Pfas*, and other reactions in purine synthesis use glutamine as a nitrogen donor.

Several groups of downregulated genes provide insight into the reduced cell growth observed at high NAM concentrations. These include downregulation of regulatory genes in growth signalling, such as WNT, MAPK and PI3K/AKT/mTOR pathway (*Sgk2*, *Tlr2*, *Angpt4*, *Il6*, *Lamc2*, *Creb3l1*, *Thbs2*, *Fgfr2*, *Dusp14, Fhl2, Socs3, Wnt4, Ccn5*), cell cycle and proliferation (*Hdgf, S100a4, Syne2, Kntc1, Spdl1*) and extracellular matrix and adhesion (*Mmp9, Mmp12, Pdpn, Col5a2, Itga5*). Growth signalling, particularly the WNT pathway, is strongly linked to aerobic glycolysis (Pate et al., 2014; Thompson, 2014; Vallée et al., 2021), and the shift from Warburg metabolism could impact these regulatory pathways to lower cell proliferation. As discussed in Sections 3.1 and 3.2, NAD^+^ dependent (and NAM inhibited) SIRTs may be impacting these regulatory processes which have been shown to interact with PI3K/AKT/mTOR pathway to reduce cell growth (Fan et al., 2023; Ghosh et al., 2010; Sadria and Layton, 2021).

Functional gene enrichment analysis using GSEA (Section 2.3 and Supplementary Table S2) further supports these metabolic changes (Figure 4b). Of the four altered pathways, three positively enriched pathways were related to oxidative phosphorylation and mitochondrial ETC activity, while the negatively enriched pathway was linked to PI3K/AKT/mTOR signalling. These findings reinforce the metabolic shift towards OXPHOS and the suppression of growth-associated signalling pathways in NAM-fed cultures.

DGE analysis was applied to NA feed supplementation. NA-fed gene expressions were much more similar to the control than NAM-fed conditions. This is evident in the PCA plot of gene expression (Supplementary Figure S3), as well as by comparing process data on day 6 in Section 3.3. Consequently, only one gene was differentially expressed, which was the downregulation of *Ogdh* (oxoglutarate dehydrogenase). *Ogdh* is a critical mitochondrial redox sensor and regulator, and its downregulation represents an early response to oxidative stress (Chang et al., 2022; McLain et al., 2011), suggesting that NA feeding may have initiated early signs of cellular stress. However, since these gene expression results were collected on day 6, prior to the significant drop in viability observed in NA-fed cultures, the later decline in cell health is not fully explained.

## 4 Conclusion

In this study, NAD⁺ precursors NAM and NA were supplemented in a fed-batch CHO cell culture. NAM supplementation reduces peak lactate by 40% and reverses the Warburg effect to induce the lactate shift three days earlier than control conditions. Transcriptomic data reveal NAM promotes a shift from glycolytic to oxidative metabolism, with an upregulation of key mitochondrial ETC genes. These positive impacts are theorised to be caused by an increase in intracellular NAD⁺ concentrations due to synthesis from NAM in the salvage pathway.

NAM supplementation presents a widely applicable and low-cost strategy to alleviate lactate accumulation, leading to several positive impacts in upstream bioprocessing. Firstly, adding NAM into a feed formulation would counter any rise in lactate levels, preventing the negative impacts of high lactate concentration on cell viability and product quality. Secondly, NAM promotes OXPHOS and suppresses cell growth, resulting in a more energy - and nutrient-efficient metabolic state. In perfusion cultures, where media consumption is a major cost driver, limiting cell proliferation reduces unnecessary biomass accumulation and nutrient waste. This allows a greater proportion of resources to be directed toward recombinant protein production, improving overall process efficiency. Thirdly, preventing lactate secretion using NAM would significantly reduce the requirement for base addition to maintain the bioreactor pH setpoint, therefore preventing hyperosmotic conditions from high base addition that may negatively affect cell culture. This would increase the capacity for enriched feeds during cell culture without causing hyperosmotic conditions, enabling the culture to achieve higher titres. This is particularly beneficial in intensified fed-batch conditions, where there is a significant demand for nutrients but also a heightened risk of hyperosmotic stress and runaway lactate accumulation.

NAM could be readily incorporated into bioreactor feeds from the beginning of process development or could be applied if development work shows a cell line has a propensity for lactate accumulation that was missed in early screening. It could also be applied as a process control response to runaway lactate accumulation during large scale production and prevent cultures. As mentioned above, NAM would be greatly beneficial to perfusion and intensified fed batch processes. However, it’s important to note that this strategy has so far only been demonstrated in a single cell line. While the underlying principles, such as redox balancing and metabolic efficiency, are broadly relevant across cell types, further validation is needed to confirm its applicability to other CHO variants or production platforms.

High concentrations of NAM attenuate cell growth with DGE and functional enrichment revealing a downregulation of growth regulatory genes, particularly in the PI3K/AKT/mTOR pathway. This is likely due to changes to the activity of NAD^+^ dependent SIRTs, decreased glycolytic fluxes as well as increased energetic and nucleoside demands from NAD⁺ synthesis. Despite this, cell-specific productivity remains unaffected, indicating that controlled NAM feeding can optimise lactate metabolism without compromising recombinant protein production. In contrast, NA supplementation failed to produce similar metabolic benefits and instead triggered a late-stage accumulation of lactate, coinciding with reduced cell viability. The downregulation of *Ogdh*, a key mitochondrial redox sensor, suggested early signs of cellular stress, potentially contributing to the later decline in culture performance.

A feed design incorporating NAM would have to carefully tune feed composition and timing to balance the positive impacts of reduced Warburg effect with the changes to cell growth and metabolism. Additional nucleosides may also need to be supplemented alongside NAM to assist the synthesis of NAD+ without draining cellular resources. Additionally, metabolomics analysis in this study revealed significantly increased consumption of glutamine and alanine (see Figure S2) with reduced glutamate, glycine and aspartate consumption. It may be beneficial to adjust the ratios of amino acids to assist with the adaptation to a high NAD+ environment when NAM is fed.

## Notes

### Competing Interest Statement

The authors have declared no competing interest.

## References

Agrawal, A., Balcı, H., Hanspers, K., Coort, S.L., Martens, M., Slenter, D.N., Ehrhart, F., Digles, D., Waagmeester, A., Wassink, I., Abbassi-Daloii, T., Lopes, E.N., Iyer, A., Acosta, J.M., Willighagen, L.G., Nishida, K., Riutta, A., Basaric, H., Evelo, C.T., Willighagen, E.L., Kutmon, M., Pico, A.R., 2024. WikiPathways 2024: next generation pathway database. Nucleic Acids Res 52. 10.1093/nar/gkad960

Alhuthali, S., Kotidis, P., Kontoravdi, C., 2021. Osmolality effects on cho cell growth, cell volume, antibody productivity and glycosylation. Int J Mol Sci 22. 10.3390/ijms22073290

Amelio, I., Cutruzzolá, F., Antonov, A., Agostini, M., Melino, G., 2014. Serine and glycine metabolism in cancer. Trends Biochem Sci. 10.1016/j.tibs.2014.02.004

Anders, S., Huber, W., 2010. Differential expression analysis for sequence count data. Genome Biol 11. 10.1186/gb-2010-11-10-r106

Audrito, V., Messana, V.G., Moiso, E., Vitale, N., Arruga, F., Brandimarte, L., Gaudino, F., Pellegrino, E., Vaisitti, T., Riganti, C., Piva, R., Deaglio, S., 2020. Nampt over-expression recapitulates the braf inhibitor resistant phenotype plasticity in melanoma. Cancers (Basel) 12. 10.3390/cancers12123855

Baldassarri, C., Giorgioni, G., Piergentili, A., Quaglia, W., Fontana, S., Mammoli, V., Minazzato, G., Marangoni, E., Gasparrini, M., Sorci, L., Raffaelli, N., Cappellacci, L., Petrelli, R., Del Bello, F., 2023. Properly Substituted Benzimidazoles as a New Promising Class of Nicotinate Phosphoribosyltransferase (NAPRT) Modulators. Pharmaceuticals 16. 10.3390/ph16020189

Becker, M., Junghans, L., Teleki, A., Bechmann, J., Takors, R., 2019. The less the better: How suppressed base addition boosts production of monoclonal antibodies with Chinese hamster ovary cells. Front Bioeng Biotechnol 7. 10.3389/fbioe.2019.00076

Bitterman, K.J., Anderson, R.M., Cohen, H.Y., Latorre-Esteves, M., Sinclair, D.A., 2002. Inhibition of silencing and accelerated aging by nicotinamide, a putative negative regulator of yeast Sir2 and human SIRT1. Journal of Biological Chemistry 277. 10.1074/jbc.M205670200

Bongarzone, S., Barbon, E., Ferocino, A., Alsulaimani, L., Dunn, J., Kim, J., Sunassee, K., Gee, A., 2020. Imaging niacin trafficking with positron emission tomography reveals in vivo monocarboxylate transporter distribution. Nucl Med Biol 88–89. 10.1016/j.nucmedbio.2020.07.002

Boroughs, L.K., Deberardinis, R.J., 2015. Metabolic pathways promoting cancer cell survival and growth. Nat Cell Biol. 10.1038/ncb3124

Buchsteiner, M., Quek, L.E., Gray, P., Nielsen, L.K., 2018. Improving culture performance and antibody production in CHO cell culture processes by reducing the Warburg effect. Biotechnol Bioeng 115. 10.1002/bit.26724

Burgos, E.S., Schramm, V.L., 2008. Weak coupling of ATP hydrolysis to the chemical equilibrium of human nicotinamide phosphoribosyltransferase. Biochemistry 47. 10.1021/bi801198m

Cairns, R.A., Harris, I.S., Mak, T.W., 2011. Regulation of cancer cell metabolism. Nat Rev Cancer. 10.1038/nrc2981

Cercillieux, A., Ratajczak, J., Joffraud, M., Sanchez-Garcia, J.L., Jacot, G., Zollinger, A., Métairon, S., Giroud-Gerbetant, J., Rumpler, M., Ciarlo, E., Valera-Alberni, M., Sambeat, A., Canto, C., 2022. Nicotinamide riboside kinase 1 protects against diet and age-induced pancreatic β-cell failure. Mol Metab 66. 10.1016/j.molmet.2022.101605

Chang, L.-C., Chiang, S.-K., Chen, S.-E., Hung, M.-C., 2022. Targeting 2-oxoglutarate dehydrogenase for cancer treatment. Am J Cancer Res 12.

Chen, S., Zhou, Y., Chen, Y., Gu, J., 2018. Fastp: An ultra-fast all-in-one FASTQ preprocessor, in: Bioinformatics. 10.1093/bioinformatics/bty560

Chong, W.P.K., Reddy, S.G., Yusufi, F.N.K., Lee, D.Y., Wong, N.S.C., Heng, C.K., Yap, M.G.S., Ho, Y.S., 2010. Metabolomics-driven approach for the improvement of Chinese hamster ovary cell growth: Overexpression of malate dehydrogenase II. J Biotechnol 147. 10.1016/j.jbiotec.2010.03.018

D’Amours, D., Desnoyers, S., D’Silva, I., Poirier, G.G., 1999. Poly(ADP-ribosyl)ation reactions in the regulation of nuclear functions. Biochemical Journal. 10.1042/0264-6021:3420249

Denu, J.M., 2007. Vitamins and Aging: Pathways to NAD+ Synthesis. Cell. 10.1016/j.cell.2007.04.023

Diaz-Ruiz, A., Lanasa, M., Garcia, J., Mora, H., Fan, F., Martin-Montalvo, A., Di Francesco, A., Calvo-Rubio, M., Salvador-Pascual, A., Aon, M.A., Fishbein, K.W., Pearson, K.J., Villalba, J.M., Navas, P., Bernier, M., de Cabo, R., 2018. Overexpression of CYB5R3 and NQO1, two NAD+-producing enzymes, mimics aspects of caloric restriction. Aging Cell 17. 10.1111/acel.12767

Doyen, D., Poët, M., Jarretou, G., Pisani, D.F., Tauc, M., Cougnon, M., Argentina, M., Bouret, Y., Counillon, L., 2022. Intracellular pH Control by Membrane Transport in Mammalian Cells. Insights Into the Selective Advantages of Functional Redundancy. Front Mol Biosci. 10.3389/fmolb.2022.825028

Fan, X., He, Y., Wu, G., Chen, H., Cheng, X., Zhan, Y., An, C., Chen, T., Wang, X., 2023. Sirt3 activates autophagy to prevent DOX-induced senescence by inactivating PI3K/AKT/mTOR pathway in A549 cells. Biochim Biophys Acta Mol Cell Res 1870. 10.1016/j.bbamcr.2022.119411

Fang, E.F., Scheibye-Knudsen, M., Brace, L.E., Kassahun, H., Sengupta, T., Nilsen, H., Mitchell, J.R., Croteau, D.L., Bohr, V.A., 2014. Defective mitophagy in XPA via PARP-1 hyperactivation and NAD +/SIRT1 reduction. Cell 157. 10.1016/j.cell.2014.03.026

Fang, F., Zhuang, P., Feng, X., Liu, P., Liu, D., Huang, H., Li, L., Chen, W., Liu, L., Sun, Y., Jiang, H., Ye, J., Hu, Y., 2022. NMNAT2 is downregulated in glaucomatous RGCs, and RGC-specific gene therapy rescues neurodegeneration and visual function. Molecular Therapy 30. 10.1016/j.ymthe.2022.01.035

Fang, Z., Liu, X., Peltz, G., 2023. GSEApy: a comprehensive package for performing gene set enrichment analysis in Python. Bioinformatics 39. 10.1093/bioinformatics/btac757

Fouquerel, E., Goellner, E.M., Yu, Z., Gagné, J.P., de Moura, M.B., Feinstein, T., Wheeler, D., Redpath, P., Li, J., Romero, G., Migaud, M., Van Houten, B., Poirier, G.G., Sobol, R.W., 2014. ARTD1/PARP1 negatively regulates glycolysis by inhibiting hexokinase 1 independent of NAD+ depletion. Cell Rep 8. 10.1016/j.celrep.2014.08.036

Fu, T., Zhang, C., Jing, Y., Jiang, C., Li, Z., Wang, S., Ma, K., Zhang, D., Hou, S., Dai, J., Kou, G., Wang, H., 2016. Regulation of cell growth and apoptosis through lactate dehydrogenase C over-expression in Chinese hamster ovary cells. Appl Microbiol Biotechnol 100. 10.1007/s00253-016-7348-4

Gardell, S.J., Hopf, M., Khan, A., Dispagna, M., Hampton Sessions, E., Falter, R., Kapoor, N., Brooks, J., Culver, J., Petucci, C., Ma, C.T., Cohen, S.E., Tanaka, J., Burgos, E.S., Hirschi, J.S., Smith, S.R., Sergienko, E., Pinkerton, A.B., 2019. Boosting NAD+ with a small molecule that activates NAMPT. Nat Commun 10. 10.1038/s41467-019-11078-z

Geueke, B., Riebel, B., Hummel, W., 2003. NADH oxidase from Lactobacillus brevis: A new catalyst for the regeneration of NAD. Enzyme Microb Technol 32. 10.1016/S0141-0229(02)00290-9

Ghorbaniaghdam, A., Chen, J., Henry, O., Jolicoeur, M., 2014. Analyzing clonal variation of monoclonal antibody-producing CHO cell lines using an in silico metabolomic platform. PLoS One 9. 10.1371/journal.pone.0090832

Ghosh, H.S., McBurney, M., Robbins, P.D., 2010. SIRT1 negatively regulates the mammalian target of rapamycin. PLoS One 5. 10.1371/journal.pone.0009199

Gopal, E., Fei, Y.J., Miyauchi, S., Zhuang, L., Prasad, P.D., Ganapathy, V., 2005. Sodium-coupled and electrogenic transport of B-complex vitamin nicotinic acid by slc5a8, a member of the Na/glucose co-transporter gene family. Biochemical Journal 388. 10.1042/BJ20041916

Gopal, E., Miyauchi, S., Martin, P.M., Ananth, S., Roon, P., Smith, S.B., Ganapathy, V., 2007. Transport of nicotinate and structurally related compounds by human SMCT1 (SLC5A8) and its relevance to drug transport in the mammalian intestinal tract. Pharm Res 24. 10.1007/s11095-006-9176-1

Greer, S.N., Metcalf, J.L., Wang, Y., Ohh, M., 2012. The updated biology of hypoxia-inducible factor. EMBO Journal. 10.1038/emboj.2012.125

Han, H.-J., Kim, H., Yu, H.G., Park, J.U., Bae, J.H., Lee, J.H., Hong, J.K., Baik, J.Y., 2024. Evaluation of NAD+ precursors for improved metabolism and productivity of antibody-producing CHO cell. Biotechnol J 19, 2400311. 10.1002/biot.202400311

Hartley, F., Walker, T., Chung, V., Morten, K., 2018. Mechanisms driving the lactate switch in Chinese hamster ovary cells. Biotechnol Bioeng. 10.1002/bit.26603

Hashida, S.N., Itami, T., Takahara, K., Hirabayashi, T., Uchimiya, H., Kawai-Yamada, M., 2016. Increased rate of NAD metabolism shortens plant longevity by accelerating developmental senescence in Arabidopsis. Plant Cell Physiol 57. 10.1093/pcp/pcw155

Hefzi, H., Martínez-Monge, I., Marin de Mas, I., Cowie, N.L., Toledo, A.G., Noh, S.M., Karottki, K.J. la C., Decker, M., Arnsdorf, J., Camacho-Zaragoza, J.M., Kol, S., Schoffelen, S., Pristovšek, N., Hansen, A.H., Miguez, A.A., Bjørn, S.P., Brøndum, K.K., Javidi, E.M., Jensen, K.L., Stangl, L., Kreidl, E., Kallehauge, T.B., Ley, D., Ménard, P., Petersen, H.M., Sukhova, Z., Bauer, A., Casanova, E., Barron, N., Malmström, J., Nielsen, L.K., Lee, G.M., Kildegaard, H.F., Voldborg, B.G., Lewis, N.E., 2025. Multiplex genome editing eliminates lactate production without impacting growth rate in mammalian cells. Nat Metab 7, 212–227. 10.1038/s42255-024-01193-7

Heiden, M.G.V., Cantley, L.C., Thompson, C.B., 2009. Understanding the warburg effect: The metabolic requirements of cell proliferation. Science (1979). 10.1126/science.1160809

Horváthová, J., Moravčík, R., Matúšková, M., Šišovský, V., Boháč, A., Zeman, M., 2021. Inhibition of glycolysis suppresses cell proliferation and tumor progression in vivo: Perspectives for chronotherapy. Int J Mol Sci 22. 10.3390/ijms22094390

Howe, K.L., Achuthan, P., Allen, James, Allen, Jamie, Alvarez-Jarreta, J., Ridwan Amode, M., Armean, I.M., Azov, A.G., Bennett, R., Bhai, J., Billis, K., Boddu, S., Charkhchi, M., Cummins, C., da Rin Fioretto, L., Davidson, C., Dodiya, K., El Houdaigui, B., Fatima, R., Gall, A., Giron, C.G., Grego, T., Guijarro-Clarke, C., Haggerty, L., Hemrom, A., Hourlier, T., Izuogu, O.G., Juettemann, T., Kaikala, V., Kay, M., Lavidas, I., Le, T., Lemos, D., Martinez, J.G., Marugán, J.C., Maurel, T., McMahon, A.C., Mohanan, S., Moore, B., Muffato, M., Oheh, D.N., Paraschas, D., Parker, A., Parton, A., Prosovetskaia, I., Sakthivel, M.P., Abdul Salam, A.I., Schmitt, B.M., Schuilenburg, H., Sheppard, D., Steed, E., Szpak, M., Szuba, M., Taylor, K., Thormann, A., Threadgold, G., Walts, B., Winterbottom, A., Chakiachvili, M., Chaubal, A., de Silva, N., Flint, B., Frankish, A., Hunt, S.E., IIsley, G.R., Langridge, N., Loveland, J.E., Martin, F.J., Mudge, J.M., Morales, J., Perry, E., Ruffier, M., Tate, J., Thybert, D., Trevanion, S.J., Cunningham, F., Yates, A.D., Zerbino, D.R., Flicek, P., 2021. Ensembl 2021. Nucleic Acids Res 49. 10.1093/nar/gkaa942

Imai, S.I., Armstrong, C.M., Kaeberlein, M., Guarente, L., 2000. Transcriptional silencing and longevity protein Sir2 is an NAD-dependent histone deacetylase. Nature 403. 10.1038/35001622

Jones, W., Bianchi, K., 2015. Aerobic glycolysis: Beyond proliferation. Front Immunol. 10.3389/fimmu.2015.00227

Jubrias, S.A., Crowther, G.J., Shankland, E.G., Gronka, R.K., Conley, K.E., 2003. Acidosis inhibits oxidative phosphorylation in contracting human skeletal muscle in vivo. Journal of Physiology 553. 10.1113/jphysiol.2003.045872

Kang, M., Park, S., Park, S.H., Lee, H.G., Park, J.H., 2022. A Double-Edged Sword: The Two Faces of PARylation. Int J Mol Sci. 10.3390/ijms23179826

Kim, H.J., Oh, G.S., Shen, A., Lee, S.B., Choe, S.K., Kwon, K.B., Lee, S., Seo, K.S., Kwak, T.H., Park, R., So, H.S., 2014. Augmentation of NAD+ by NQO1 attenuates cisplatin-mediated hearing impairment. Cell Death Dis 5. 10.1038/cddis.2014.255

Kunert, R., Reinhart, D., 2016. Advances in recombinant antibody manufacturing. Appl Microbiol Biotechnol. 10.1007/s00253-016-7388-9

Lao, M.S., Toth, D., 1997. Effects of ammonium and lactate on growth and metabolism of a recombinant Chinese hamster ovary cell culture. Biotechnol Prog 13. 10.1021/bp9602360

Lee, J.H., Kang, H.I., Kim, S., Ahn, Y. Bin, Kim, H., Hong, J.K., Baik, J.Y., 2023. NAD+ supplementation improves mAb productivity in CHO cells via a glucose metabolic shift. Biotechnol J 18. 10.1002/biot.202200570

Lee, M.S., Kim, K.W., Kim, Y.H., Lee, G.M., 2003. Proteome Analysis of Antibody-Expressing CHO Cells in Response to Hyperosmotic Pressure. Biotechnol Prog 19. 10.1021/bp034093a

Li, J., Wong, C.L., Vijayasankaran, N., Hudson, T., Amanullah, A., 2012. Feeding lactate for CHO cell culture processes: Impact on culture metabolism and performance. Biotechnol Bioeng 109. 10.1002/bit.24389

Liang, J., Wang, P., Wei, J., Bao, C., Han, D., 2015. Nicotinamide Mononucleotide Adenylyltransferase 1 Protects Neural Cells Against Ischemic Injury in Primary Cultured Neuronal Cells and Mouse Brain with Ischemic Stroke Through AMP-Activated Protein Kinase Activation. Neurochem Res 40. 10.1007/s11064-015-1569-2

Liberti, M. V., Locasale, J.W., 2016. The Warburg Effect: How Does it Benefit Cancer Cells? Trends Biochem Sci. 10.1016/j.tibs.2015.12.001

Liu, L., Su, X., Quinn, W.J., Hui, S., Krukenberg, K., Frederick, D.W., Redpath, P., Zhan, L., Chellappa, K., White, E., Migaud, M., Mitchison, T.J., Baur, J.A., Rabinowitz, J.D., 2018. Quantitative Analysis of NAD Synthesis-Breakdown Fluxes. Cell Metab 27. 10.1016/j.cmet.2018.03.018

Luengo, A., Li, Z., Gui, D.Y., Sullivan, L.B., Zagorulya, M., Do, B.T., Ferreira, R., Naamati, A., Ali, A., Lewis, C.A., Thomas, C.J., Spranger, S., Matheson, N.J., Vander Heiden, M.G., 2021. Increased demand for NAD+ relative to ATP drives aerobic glycolysis. Mol Cell 81. 10.1016/j.molcel.2020.12.012

Lunt, S.Y., Vander Heiden, M.G., 2011. Aerobic glycolysis: Meeting the metabolic requirements of cell proliferation. Annu Rev Cell Dev Biol 27. 10.1146/annurev-cellbio-092910-154237

McLain, A.L., Szweda, P.A., Szweda, L.I., 2011. α-Ketoglutarate dehydrogenase: A mitochondrial redox sensor. Free Radic Res. 10.3109/10715762.2010.534163

Mulukutla, B.C., Gramer, M., Hu, W.S., 2012. On metabolic shift to lactate consumption in fed-batch culture of mammalian cells. Metab Eng 14. 10.1016/j.ymben.2011.12.006

North, B.J., Verdin, E., 2004. Sirtuins: Sir2-related NAD-dependent protein deacetylases. Genome Biol. 10.1186/gb-2004-5-5-224

Pan, X., Alsayyari, A.A., Dalm, C., Hageman, J.A., Wijffels, R.H., Martens, D.E., 2019. Transcriptome Analysis of CHO Cell Size Increase During a Fed-Batch Process. Biotechnol J 14. 10.1002/biot.201800156

Pate, K.T., Stringari, C., Sprowl-Tanio, S., Wang, K., TeSlaa, T., Hoverter, N.P., McQuade, M.M., Garner, C., Digman, M.A., Teitell, M.A., Edwards, R.A., Gratton, E., Waterman, M.L., 2014. Wnt signaling directs a metabolic program of glycolysis and angiogenesis in colon cancer. EMBO J 33. 10.15252/embj.201488598

Patro, R., Duggal, G., Love, M.I., Irizarry, R.A., Kingsford, C., 2017. Salmon provides fast and bias-aware quantification of transcript expression. Nat Methods 14. 10.1038/nmeth.4197

Pedregosa, F., Varoquaux, G., Gramfort, A., Michel, V., Thirion, B., Grisel, O., Blondel, M., Prettenhofer, P., Weiss, R., Dubourg, V., Vanderplas, J., Passos, A., Cournapeau, D., Brucher, M., Perrot, M., Duchesnay, É., 2011. Scikit-learn: Machine learning in Python. Journal of Machine Learning Research 12.

Pinho, A. V., Mawson, A., Gill, A., Arshi, M., Warmerdam, M., Giry-Laterriere, M., Eling, N., Lie, T., Kuster, E., Camargo, S., Biankin, A. V., Wu, J., Rooman, I., 2016. Sirtuin1 stimulates the proliferation and the expression of glycolysis genes in pancreatic neoplastic lesions. Oncotarget 7. 10.18632/oncotarget.11013

Pirinen, E., Cantó, C., Jo, Y.S., Morato, L., Zhang, H., Menzies, K.J., Williams, E.G., Mouchiroud, L., Moullan, N., Hagberg, C., Li, W., Timmers, S., Imhof, R., Verbeek, J., Pujol, A., Van Loon, B., Viscomi, C., Zeviani, M., Schrauwen, P., Sauve, A.A., Schoonjans, K., Auwerx, J., 2014. Pharmacological inhibition of poly(ADP-ribose) polymerases improves fitness and mitochondrial function in skeletal muscle. Cell Metab 19. 10.1016/j.cmet.2014.04.002

Poburko, D., Santo-Domingo, J., Demaurex, N., 2011. Dynamic regulation of the mitochondrial proton gradient during cytosolic calcium elevations. Journal of Biological Chemistry 286. 10.1074/jbc.M110.159962

Price, N.L., Gomes, A.P., Ling, A.J.Y., Duarte, F. V., Martin-Montalvo, A., North, B.J., Agarwal, B., Ye, L., Ramadori, G., Teodoro, J.S., Hubbard, B.P., Varela, A.T., Davis, J.G., Varamini, B., Hafner, A., Moaddel, R., Rolo, A.P., Coppari, R., Palmeira, C.M., De Cabo, R., Baur, J.A., Sinclair, D.A., 2012. SIRT1 is required for AMPK activation and the beneficial effects of resveratrol on mitochondrial function. Cell Metab 15. 10.1016/j.cmet.2012.04.003

Rodgers, J.T., Lerin, C., Haas, W., Gygi, S.P., Spiegelman, B.M., Puigserver, P., 2005. Nutrient control of glucose homeostasis through a complex of PGC-1α and SIRT1. Nature 434. 10.1038/nature03354

Romanova, N., Schelletter, L., Hoffrogge, R., Noll, T., 2022. Hyperosmolality in CHO cell culture: effects on the proteome. Appl Microbiol Biotechnol 106. 10.1007/s00253-022-11861-x

Rossi, F., Geiszler, P.C., Meng, W., Barron, M.R., Prior, M., Herd-Smith, A., Loreto, A., Lopez, M.Y., Faas, H., Pardon, M.C., Conforti, L., 2018. NAD-biosynthetic enzyme NMNAT1 reduces early behavioral impairment in the htau mouse model of tauopathy. Behavioural Brain Research 339. 10.1016/j.bbr.2017.11.030

Sacco, S.A., McAtee Pereira, A.G., Trenary, I., Smith, K.D., Betenbaugh, M.J., Young, J.D., 2023. Overexpression of peroxisome proliferator-activated receptor γ co-activator-1⍺ (PGC-1⍺) in Chinese hamster ovary cells increases oxidative metabolism and IgG productivity. Metab Eng 79. 10.1016/j.ymben.2023.07.005

Sadria, M., Layton, A.T., 2021. Interactions among mTORC, AMPK and SIRT: a computational model for cell energy balance and metabolism. Cell Communication and Signaling 19. 10.1186/s12964-021-00706-1

Sánchez, B.J., Zhang, C., Nilsson, A., Lahtvee, P., Kerkhoven, E.J., Nielsen, J., 2017. Improving the phenotype predictions of a yeast genome-scale metabolic model by incorporating enzymatic constraints. Mol Syst Biol 13, 935. 10.15252/msb.20167411

Sarkar, A., Dutta, S., Sur, M., Chakraborty, S., Dey, P., Mukherjee, P., 2023. Early loss of endogenous NAD+ following rotenone treatment leads to mitochondrial dysfunction and Sarm1 induction that is ameliorated by PARP inhibition. FEBS Journal 290. 10.1111/febs.16652

Selivanov, V.A., Zeak, J.A., Roca, J., Cascante, M., Trucco, M., Votyakova, T. V., 2008. The role of external and matrix pH in mitochondrial reactive oxygen species generation. Journal of Biological Chemistry 283. 10.1074/jbc.M801019200

Soneson, C., Love, M.I., Robinson, M.D., 2016. Differential analyses for RNA-seq: Transcript-level estimates improve gene-level inferences. F1000Res 4. 10.12688/F1000RESEARCH.7563.2

Srivastava, A., Malik, L., Sarkar, H., Zakeri, M., Almodaresi, F., Soneson, C., Love, M.I., Kingsford, C., Patro, R., 2020. Alignment and mapping methodology influence transcript abundance estimation. Genome Biol 21. 10.1186/s13059-020-02151-8

Subramanian, A., Kuehn, H., Gould, J., Tamayo, P., Mesirov, J.P., 2007. GSEA-P: A desktop application for gene set enrichment analysis. Bioinformatics 23. 10.1093/bioinformatics/btm369

Takahashi, R., Nakamura, S., Nakazawa, T., Minoura, K., Yoshida, T., Nishi, Y., Kobayashi, Y., Ohkubo, T., 2010. Structure and reaction mechanism of human nicotinamide phosphoribosyltransferase. J Biochem 147. 10.1093/jb/mvp152

Talla, V., Koilkonda, R., Guy, J., 2020. Gene Therapy with Single-Subunit Yeast NADH-Ubiquinone Oxidoreductase (NDI1) Improves the Visual Function in Experimental Autoimmune Encephalomyelitis (EAE) Mice Model of Multiple Sclerosis (MS). Mol Neurobiol 57. 10.1007/s12035-019-01857-6

Thompson, C.B., 2014. Wnt meets Warburg: another piece in the puzzle? EMBO J 33. 10.15252/embj.201488785

Titov, D. V., Cracan, V., Goodman, R.P., Peng, J., Grabarek, Z., Mootha, V.K., 2016. Complementation of mitochondrial electron transport chain by manipulation of the NAD+/NADH ratio. Science (1979) 352. 10.1126/science.aad4017

Torres, M., Altamirano, C., Dickson, A.J., 2018. Process and metabolic engineering perspectives of lactate production in mammalian cell cultures. Curr Opin Chem Eng. 10.1016/j.coche.2018.10.004

Vallée, A., Lecarpentier, Y., Vallée, J.N., 2021. The key role of the wnt/β-catenin pathway in metabolic reprogramming in cancers under normoxic conditions. Cancers (Basel). 10.3390/cancers13215557

Vemuri, G.N., Eiteman, M.A., McEwen, J.E., Olsson, L., Nielsen, J., 2007. Increasing NADH oxidation reduces overflow metabolism in Saccharomyces cerevisiae. Proc Natl Acad Sci U S A 104. 10.1073/pnas.0607469104

Wahrheit, J., Niklas, J., Heinzle, E., 2014. Metabolic control at the cytosol-mitochondria interface in different growth phases of CHO cells. Metab Eng 23. 10.1016/j.ymben.2014.02.001

Walsh, G., Walsh, E., 2022. Biopharmaceutical benchmarks 2022. Nat Biotechnol 40. 10.1038/s41587-022-01582-x

Wang, B., Hasan, M.K., Alvarado, E., Yuan, H., Wu, H., Chen, W.Y., 2011. NAMPT overexpression in prostate cancer and its contribution to tumor cell survival and stress response. Oncogene 30. 10.1038/onc.2010.468

Xie, N., Zhang, L., Gao, W., Huang, C., Huber, P.E., Zhou, X., Li, C., Shen, G., Zou, B., 2020. NAD+ metabolism: pathophysiologic mechanisms and therapeutic potential. Signal Transduct Target Ther. 10.1038/s41392-020-00311-7

Yanase, H., Takebe, K., Nio-Kobayashi, J., Takahashi-Iwanaga, H., Iwanaga, T., 2008. Cellular expression of a sodium-dependent monocarboxylate transporter (Slc5a8) and the MCT family in the mouse kidney. Histochem Cell Biol 130. 10.1007/s00418-008-0490-z

Yao, H., Liu, M., Wang, L., Zu, Y., Wu, C., Li, C., Zhang, R., Lu, H., Li, F., Xi, S., Chen, S., Gu, X., Liu, T., Cai, J., Wang, S., Yang, M., Xing, G.G., Xiong, W., Hua, L., Tang, Y., Wang, G., 2022. Discovery of small-molecule activators of nicotinamide phosphoribosyltransferase (NAMPT) and their preclinical neuroprotective activity. Cell Res 32. 10.1038/s41422-022-00651-9

Yeo, H.C., Hong, J., Lakshmanan, M., Lee, D.Y., 2020. Enzyme capacity-based genome scale modelling of CHO cells. Metab Eng 60, 138–147. 10.1016/j.ymben.2020.04.005

Zalai, D., Koczka, K., Párta, L., Wechselberger, P., Klein, T., Herwig, C., 2015. Combining mechanistic and data-driven approaches to gain process knowledge on the control of the metabolic shift to lactate uptake in a fed-batch CHO process. Biotechnol Prog 31. 10.1002/btpr.2179

Zheng, J., 2012. Energy metabolism of cancer: Glycolysis versus oxidative phosphorylation (review). Oncol Lett. 10.3892/ol.2012.928

Zhu, M.M., Goyal, A., Rank, D.L., Gupta, S.K., Vanden Boom, T., Lee, S.S., 2005. Effects of elevated pCO2 and osmolality on growth of CHO cells and production of antibody-fusion protein B1: A case study. Biotechnol Prog 21. 10.1021/bp049815s

