## Supplementary Document for "Nicotinamide reverses the Warburg effect in Chinese hamster ovary cell culture"

#### Figure S1 Recombinant product titre

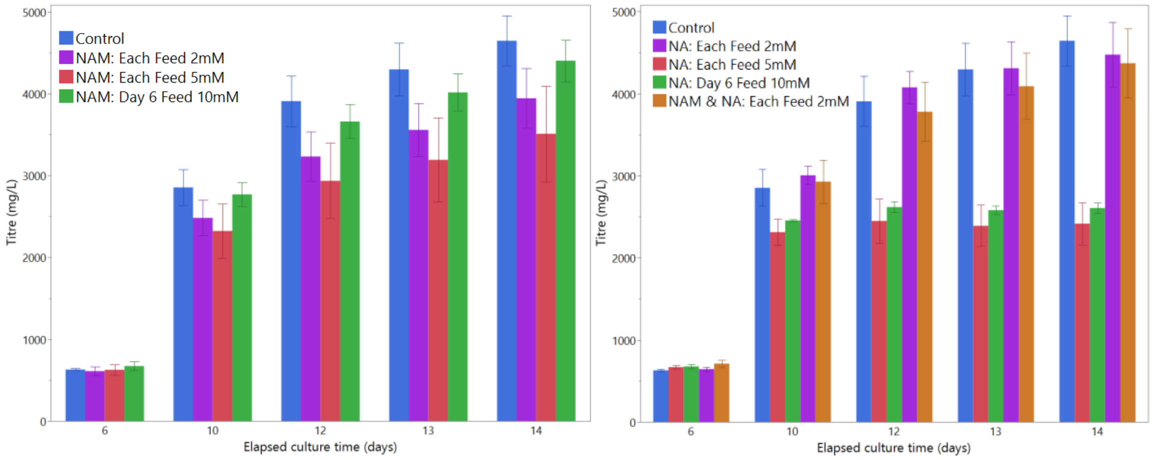


*Figure S1: Recombinant product titre for a) nicotinamide (NAM) fed cultures b) nicotinic acid (NA) fed cultures. Error bars represent standard error from the triplicates.*

#### Figure S2 Amino acid concentration profiles (normalised)

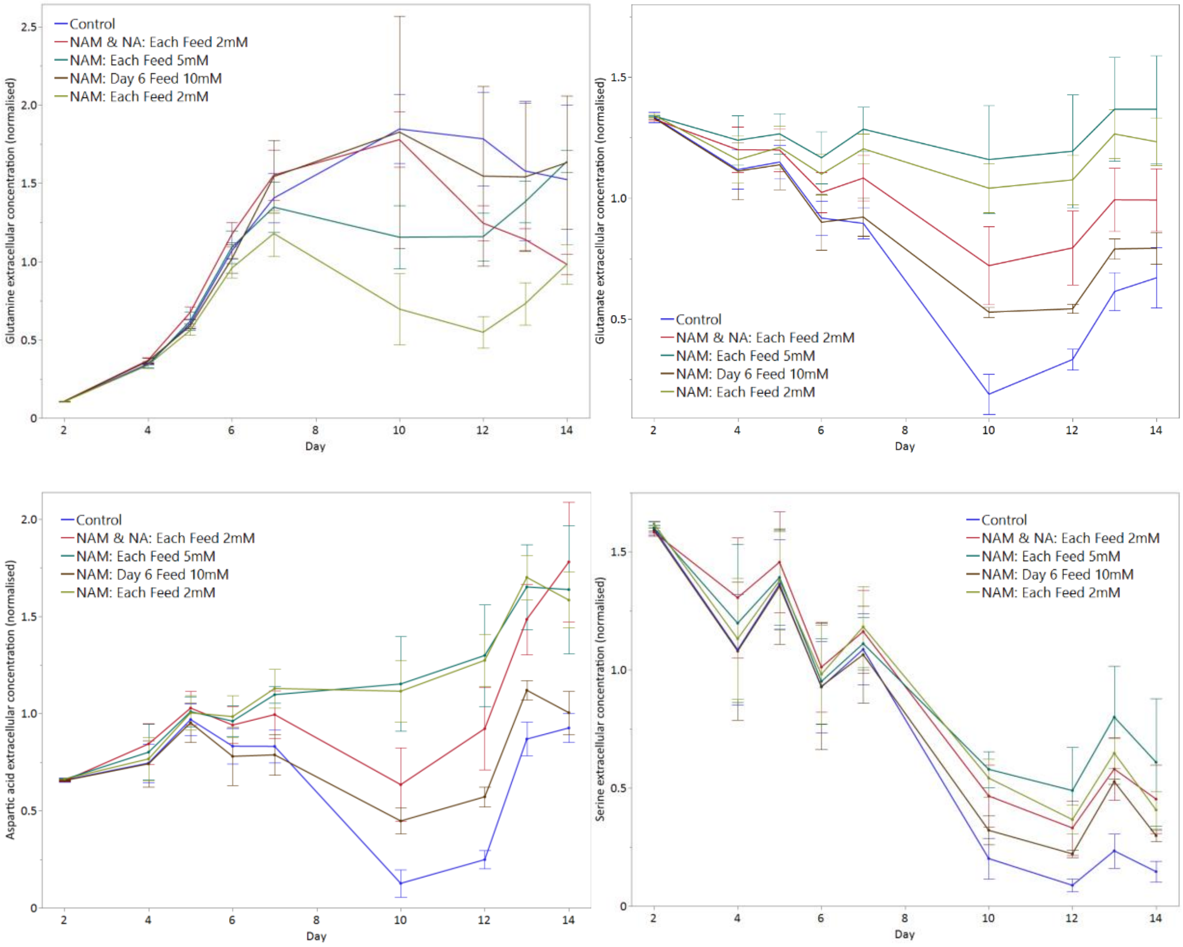

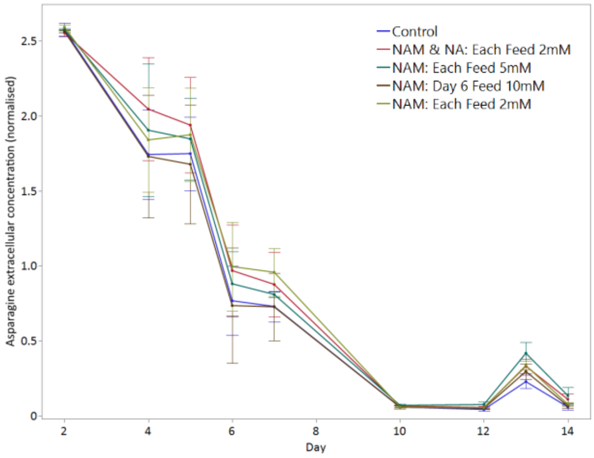
Control
NAM & NA: Each Feed 2mM
NAM: Each Feed 5mM
NAM: Day 6 Feed 10mM
NAM: Each Feed 2mM
Asparagine extracellular concentration (normalised)
Day

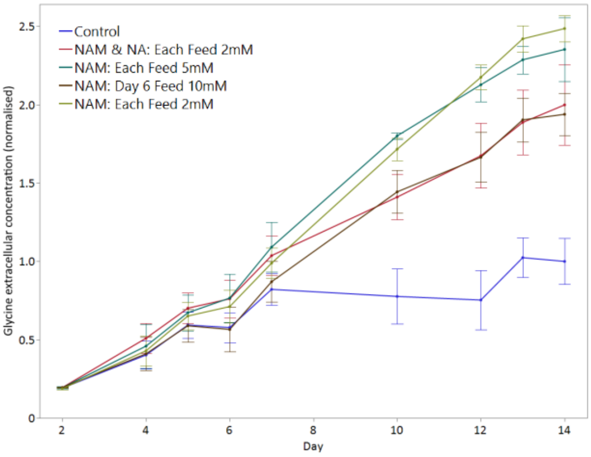
Control
NAM & NA: Each Feed 2mM
NAM: Each Feed 5mM
NAM: Day 6 Feed 10mM
NAM: Each Feed 2mM
Glycine extracellular concentration (normalised)
Day

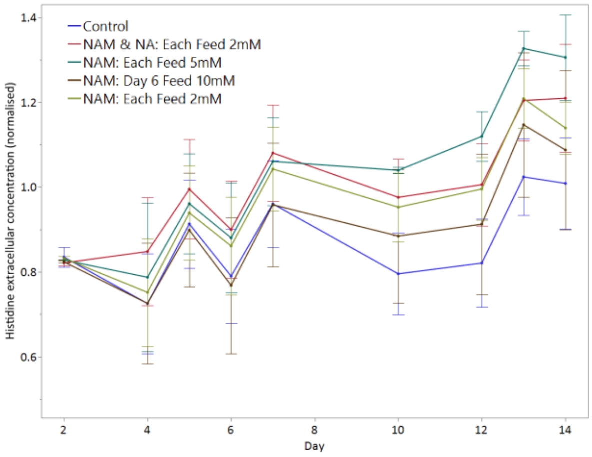
Control
NAM & NA: Each Feed 2mM
NAM: Each Feed 5mM
NAM: Day 6 Feed 10mM
NAM: Each Feed 2mM
Histidine extracellular concentration (normalised)
Day

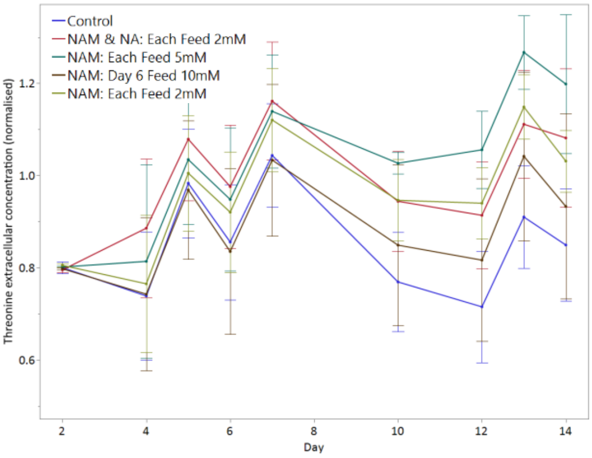
Control
NAM & NA: Each Feed 2mM
NAM: Each Feed 5mM
NAM: Day 6 Feed 10mM
NAM: Each Feed 2mM
Threonine extracellular concentration (normalised)
Day

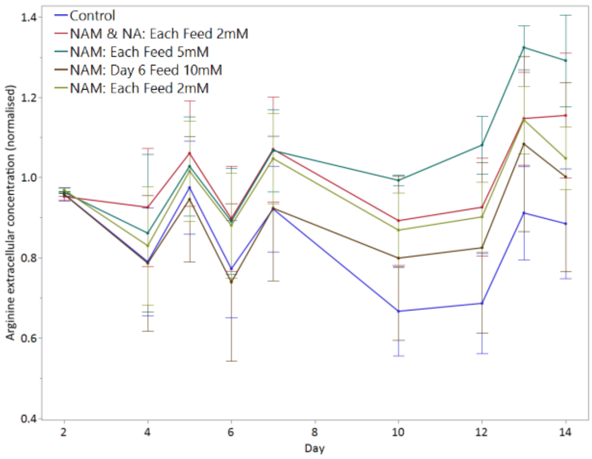
Control
NAM & NA: Each Feed 2mM
NAM: Each Feed 5mM
NAM: Day 6 Feed 10mM
NAM: Each Feed 2mM
Arginine extracellular concentration (normalised)
Day

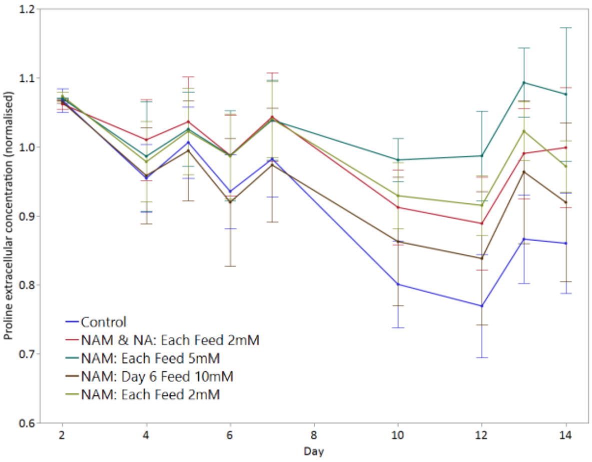
Control
NAM & NA: Each Feed 2mM
NAM: Each Feed 5mM
NAM: Day 6 Feed 10mM
NAM: Each Feed 2mM
Proline extracellular concentration (normalised)
Day

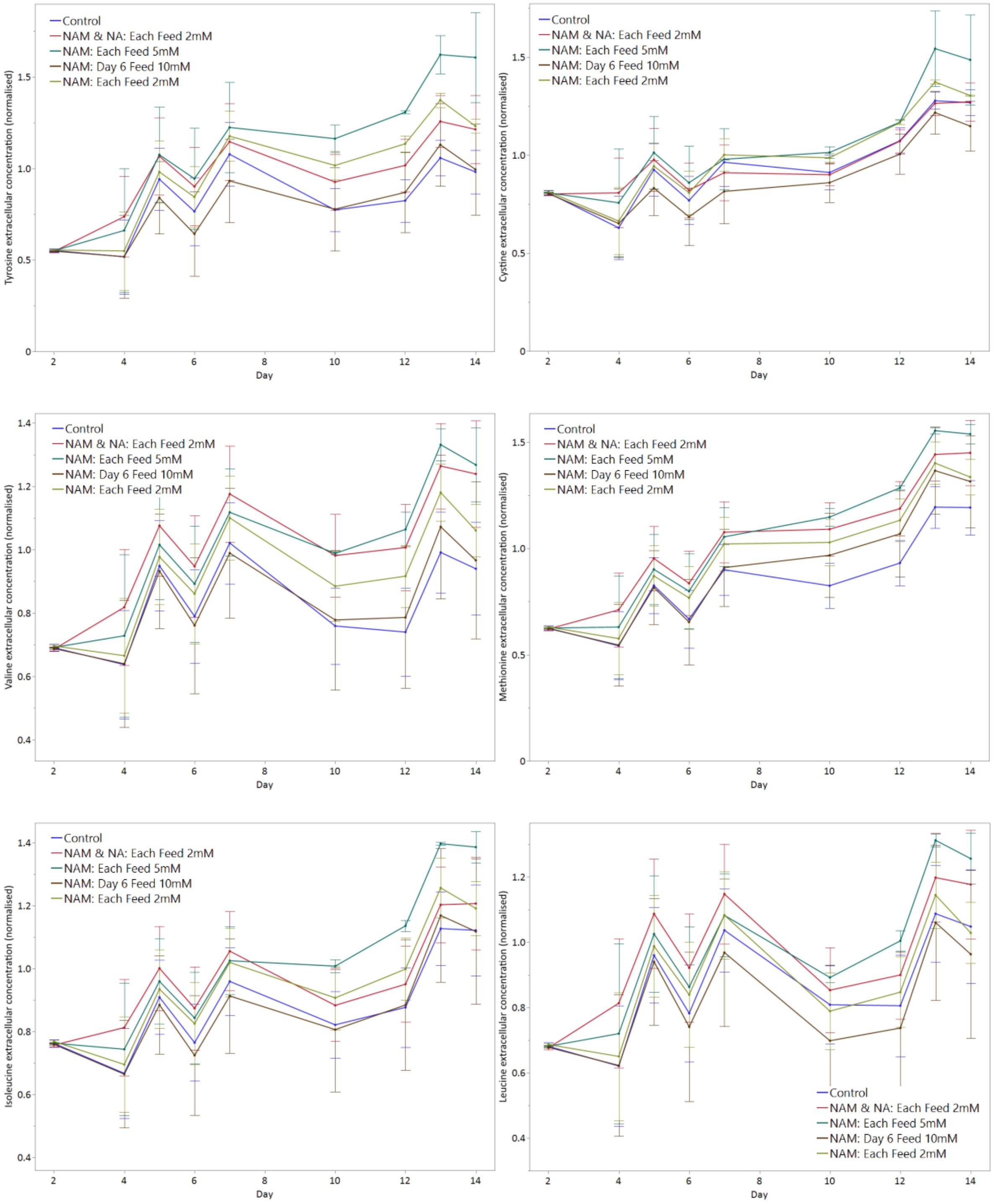
Tyrosine extracellular concentration (normalised)
Control
NAM & NA: Each Feed 2mM
NAM: Each Feed 5mM
NAM: Day 6 Feed 10mM
NAM: Each Feed 2mM
Day
Cystine extracellular concentration (normalised)
Control
NAM & NA: Each Feed 2mM
NAM: Each Feed 5mM
NAM: Day 6 Feed 10mM
NAM: Each Feed 2mM
Day
Valine extracellular concentration (normalised)
Control
NAM & NA: Each Feed 2mM
NAM: Each Feed 5mM
NAM: Day 6 Feed 10mM
NAM: Each Feed 2mM
Day
Methionine extracellular concentration (normalised)
Control
NAM & NA: Each Feed 2mM
NAM: Each Feed 5mM
NAM: Day 6 Feed 10mM
NAM: Each Feed 2mM
Day
Isoleucine extracellular concentration (normalised)
Control
NAM & NA: Each Feed 2mM
NAM: Each Feed 5mM
NAM: Day 6 Feed 10mM
NAM: Each Feed 2mM
Day
Leucine extracellular concentration (normalised)
Control
NAM & NA: Each Feed 2mM
NAM: Each Feed 5mM
NAM: Day 6 Feed 10mM
NAM: Each Feed 2mM
Day

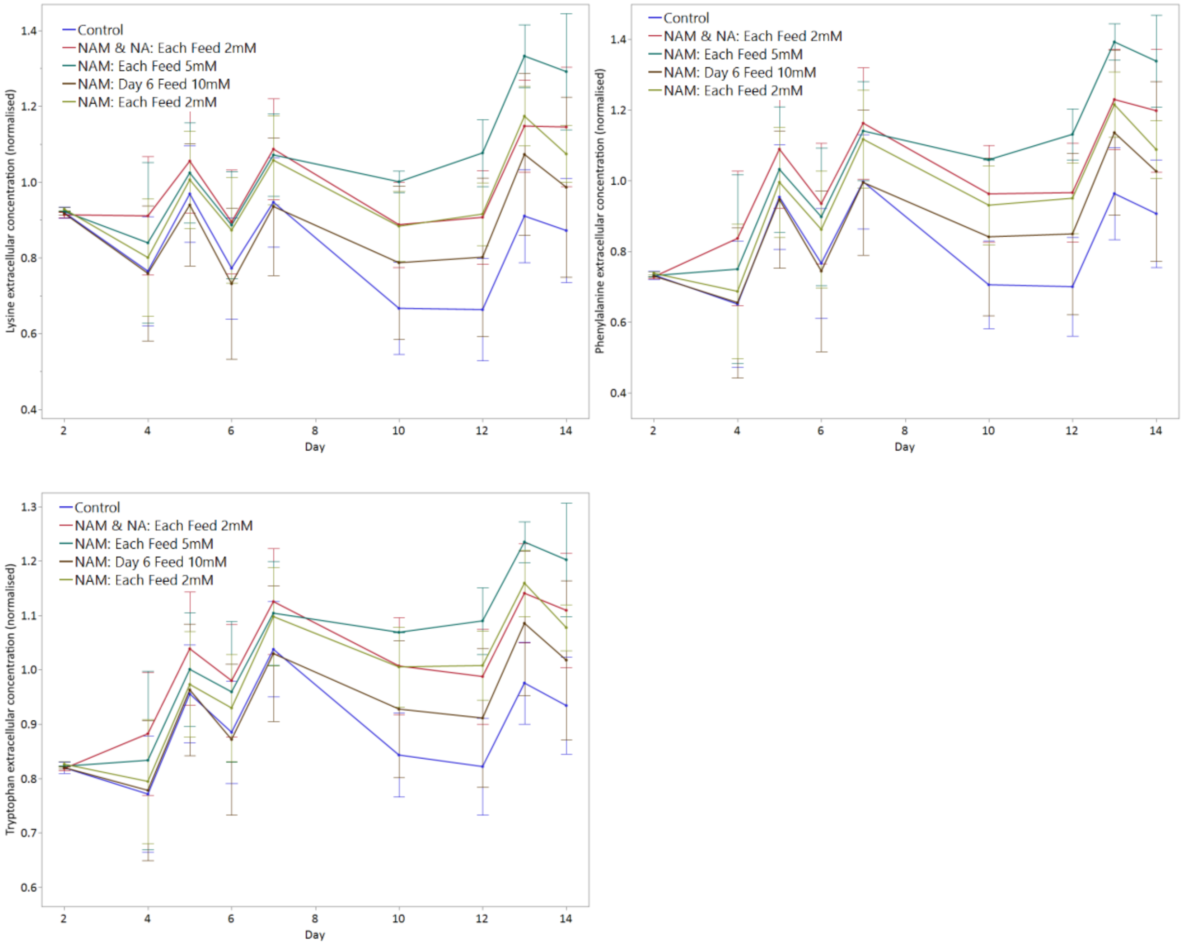


*Figure S2: Normalised amino acid concentrations for NAM supplementation experiments. Error bars represent standard error from the triplicates.*

### Figure S3 Transcriptomics PCA

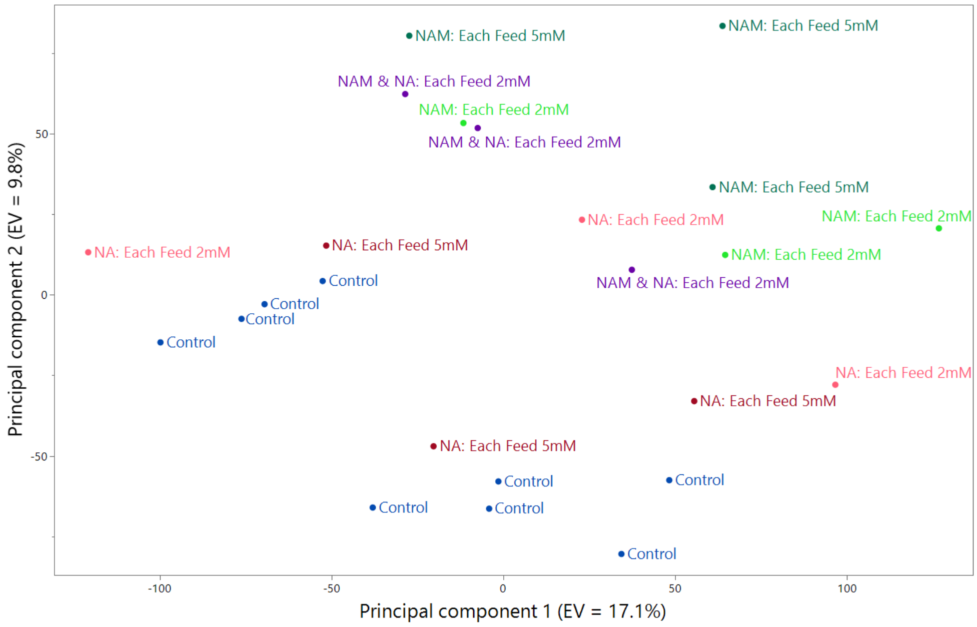


*Figure S3: Principal Component Analysis (PCA) plot showing the distribution of log2 (TPM+1) gene expression based on the first two principal components (PCs). PC1 and PC2 explain 17.1% and 9.8% of variance.*

#### Table S1 Differentially Expressed Genes

*Table S1: List of up and downregulated genes from control to NAM-fed conditions using DESeq2.*

| Ensembl_gene_id | Gene Name | Gene Description | Log2FC | $p_{adj}$ |
|---|---|---|---|---|
| ENSCGRG00001000006 | ND1 | NADH dehydrogenase subunit 1 | 0.820 | 2.9E-07 |
| ENSCGRG00001000010 | ND2 | NADH dehydrogenase subunit 2 | 0.747 | 3.7E-05 |
| ENSCGRG00001000022 | ATP6 | ATP synthase F0 subunit 6 | 0.395 | 1.4E-04 |
| ENSCGRG00001000025 | ND3 | NADH dehydrogenase subunit 3 | 0.693 | 6.2E-12 |
| ENSCGRG00001000027 | ND4L | NADH dehydrogenase subunit 4L | 0.556 | 4.7E-08 |
| ENSCGRG00001000028 | ND4 | NADH dehydrogenase subunit 4 | 0.496 | 1.9E-06 |
| ENSCGRG00001000032 | ND5 | NADH dehydrogenase subunit 5 | 0.658 | 5.8E-04 |
| ENSCGRG00001000035 | CYTB | cytochrome b | 0.572 | 2.1E-08 |
| ENSCGRG00001000139 | | serpin family B member 1 | -0.355 | 2.2E-08 |
| ENSCGRG00001000197 | Gmip | GEM interacting protein | -0.345 | 1.2E-04 |
| ENSCGRG00001000424 | Zranb3 | zinc finger RANBP2-type containing 3 | 0.322 | 2.4E-04 |
| ENSCGRG00001000692 | Lpin1 | lipin 1 | -0.315 | 3.7E-04 |
| ENSCGRG00001000968 | Itpk1 | inositol-tetrakisphosphate 1-kinase | -0.357 | 7.4E-08 |
| ENSCGRG00001001031 | Ebp | EBP cholestenol delta-isomerase | -0.341 | 2.0E-05 |
| ENSCGRG00001001177 | Sprr3 | small proline-rich protein 3 | -1.287 | 8.7E-08 |
| ENSCGRG00001001302 | | Bet1 golgi vesicular membrane trafficking protein like | 0.324 | 2.4E-04 |
| ENSCGRG00001001400 | Bmi1 | polycomb complex protein BMI-1 | 0.327 | 6.6E-12 |
| ENSCGRG00001001718 | Dxo | decapping exoribonuclease | -0.490 | 1.3E-04 |
| ENSCGRG00001001915 | Aldh3b1 | aldehyde dehydrogenase family 3 member B1 | -0.550 | 7.0E-04 |
| ENSCGRG00001002000 | Srd5a3 | steroid 5 alpha-reductase 3 | 0.398 | 1.8E-05 |
| ENSCGRG00001002061 | Hk2 | hexokinase 2 | -0.314 | 1.3E-06 |
| ENSCGRG00001002232 | Wdr77 | WD repeat domain 77 | -0.365 | 5.1E-09 |
| ENSCGRG00001002299 | Socs3 | suppressor of cytokine signaling 3 | -0.831 | 4.9E-08 |
| ENSCGRG00001002340 | Hmga1 | Cricetulus griseus high mobility group AT-hook 1 (Hmga1), mRNA. | -0.325 | 7.3E-04 |
| ENSCGRG00001002444 | Dhx29 | DExH-box helicase 29 | 0.308 | 4.6E-04 |
| ENSCGRG00001002481 | Zbtb48 | zinc finger and BTB domain containing 48 | 0.346 | 6.3E-06 |
| ENSCGRG00001002527 | St3gal5 | ST3 beta-galactoside alpha-2,3-sialyltransferase 5 | -0.308 | 2.3E-07 |
| ENSCGRG00001002618 | Thbs1 | thrombospondin 1 | 0.721 | 1.3E-05 |
| ENSCGRG00001003257 | Phkb | phosphorylase kinase beta | 0.349 | 4.8E-14 |
| ENSCGRG00001003406 | Sgk2 | serum/glucocorticoid regulated kinase 2 | -0.708 | 1.5E-04 |
| ENSCGRG00001003538 | Polr2a | polymerase (RNA) II (DNA directed) polypeptide A | 0.301 | 1.1E-06 |
| ENSCGRG00001003959 | Vnn1 | vanin 1 | -0.369 | 1.5E-20 |
| ENSCGRG00001004106 | Arid2 | AT-rich interaction domain 2 | 0.666 | 1.5E-06 |
| ENSCGRG00001004189 | Rft1 | RFT1 homolog | 0.349 | 4.6E-05 |
| ENSCGRG00001004252 | Pla1a | phospholipase A1 member A | -0.478 | 1.3E-04 |
| ENSCGRG00001004304 | Mx2 | MX dynamin like GTPase 1 | -0.326 | 2.5E-04 |
| ENSCGRG00001004544 | PEX7 | Peroxisomal targeting signal 2 receptor | 0.644 | 1.1E-12 |

| | | | | |
|---|---|---|---|---|
| ENSCGRG00001004701 | U90926 | cDNA sequence U90926 | -0.961 | 7.2E-12 |
| ENSCGRG00001005179 | Ahr | aryl-hydrocarbon receptor | -0.313 | 3.4E-05 |
| ENSCGRG00001005188 | Idi1 | isopentenyl-diphosphate delta isomerase 1 | -0.379 | 5.1E-04 |
| ENSCGRG00001005575 | Arhgap28 | Rho GTPase activating protein 28 | -0.302 | 1.1E-04 |
| ENSCGRG00001005814 | Pfas | phosphoribosylformylglycinamidine synthase | 0.334 | 1.1E-26 |
| ENSCGRG00001005835 | Dpy19l1 | dpy-19 like C-mannosyltransferase 1 | 0.306 | 1.0E-05 |
| ENSCGRG00001006775 | | | -0.540 | 7.0E-04 |
| ENSCGRG00001007148 | Arrb1 | arrestin beta 1 | -0.348 | 3.8E-05 |
| ENSCGRG00001007284 | Pclo | piccolo (presynaptic cytomatrix protein) | 0.442 | 8.5E-05 |
| ENSCGRG00001007518 | Rps6ka3 | ribosomal protein S6 kinase A3 | -0.306 | 2.1E-11 |
| ENSCGRG00001008116 | Coro7 | coronin 7 | 0.321 | 1.4E-04 |
| ENSCGRG00001008128 | Abca2 | ATP binding cassette subfamily A member 2 | 0.319 | 7.3E-14 |
| ENSCGRG00001008336 | Flrt3 | fibronectin leucine rich transmembrane protein 3 | -0.369 | 7.3E-04 |
| ENSCGRG00001008433 | Sprr1a | cornifin alpha | -0.418 | 4.0E-13 |
| ENSCGRG00001008708 | Pkib | cAMP-dependent protein kinase inhibitor beta | -0.587 | 4.1E-04 |
| ENSCGRG00001008950 | Rnaseh1 | ribonuclease H1 | 0.309 | 6.5E-05 |
| ENSCGRG00001009260 | Lig1 | DNA ligase 1 | 0.419 | 1.2E-10 |
| ENSCGRG00001009394 | Dennd3 | DENN/MADD domain containing 3 | -0.391 | 2.9E-05 |
| ENSCGRG00001009471 | Plin4 | perilipin 4 | -0.402 | 2.0E-05 |
| ENSCGRG00001009536 | Mmp19 | matrix metallopeptidase 19 | -0.422 | 2.7E-19 |
| ENSCGRG00001009756 | Gpd1 | glycerol-3-phosphate dehydrogenase 1 | -1.440 | 1.2E-06 |
| ENSCGRG00001009763 | Plac1 | placenta enriched 1 | -0.822 | 4.5E-18 |
| ENSCGRG00001009791 | Rhou | ras homolog family member U | -0.356 | 5.3E-04 |
| ENSCGRG00001009843 | Wdr25 | WD repeat domain 25 | 0.399 | 9.0E-04 |
| ENSCGRG00001009874 | Pnpla7 | patatin-like phospholipase domain containing 7 | 0.319 | 5.6E-05 |
| ENSCGRG00001010017 | Neu1 | neuraminidase 1 | -0.421 | 3.4E-08 |
| ENSCGRG00001010127 | Dusp14 | dual specificity phosphatase 14 | -0.375 | 2.5E-04 |
| ENSCGRG00001010161 | Ivns1abp | influenza virus NS1A binding protein | -0.344 | 1.4E-04 |
| ENSCGRG00001010180 | Lamc2 | laminin, gamma 2 | -0.545 | 3.5E-08 |
| ENSCGRG00001010295 | Slc39a3 | solute carrier family 39 member 3 | 0.335 | 3.7E-05 |
| ENSCGRG00001010623 | Nucb2 | nucleobindin 2 | -0.333 | 1.2E-12 |
| ENSCGRG00001010637 | Cgnl1 | cingulin like 1 | -0.693 | 7.1E-04 |
| ENSCGRG00001010709 | Fancd2 | FA complementation group D2 | 0.470 | 3.1E-11 |
| ENSCGRG00001010778 | Rtl6 | retrotransposon Gag like 6 | 0.427 | 8.4E-04 |
| ENSCGRG00001010805 | Car5b | carbonic anhydrase 5B | -0.360 | 5.4E-04 |
| ENSCGRG00001010830 | Eps8 | epidermal growth factor receptor pathway substrate 8 | -0.334 | 6.0E-06 |
| ENSCGRG00001010926 | Zfp36l1 | ZFP36 ring finger protein like 1 | -0.323 | 5.6E-07 |
| ENSCGRG00001011026 | Cidec | cell death inducing DFFA like effector c | -0.493 | 9.5E-15 |
| ENSCGRG00001011030 | | | -0.477 | 1.7E-05 |

| | | | | |
|---|---|---|---|---|
| ENSCGRG00001011164 | Amh | anti-Mullerian hormone | -0.887 | 6.1E-08 |
| ENSCGRG00001011237 | Pmvk | phosphomevalonate kinase | -0.393 | 5.3E-04 |
| ENSCGRG00001011354 | Krt80 | keratin 80 | -0.359 | 2.9E-07 |
| ENSCGRG00001011406 | Cep250 | centrosomal protein 250 | 0.308 | 1.5E-05 |
| ENSCGRG00001011598 | Creb3l1 | cAMP responsive element binding protein 3 like 1 | -0.448 | 2.1E-15 |
| ENSCGRG00001011621 | Dtwd1 | DTW domain containing 1 | 0.320 | 2.8E-10 |
| ENSCGRG00001011698 | Sfpq | splicing factor proline and glutamine rich | 0.340 | 3.0E-04 |
| ENSCGRG00001011714 | Abcb10 | ATP binding cassette subfamily B member 10 | 0.330 | 2.5E-04 |
| ENSCGRG00001011822 | Scai | suppressor of cancer cell invasion | 0.434 | 1.4E-04 |
| ENSCGRG00001011882 | Prss35 | serine protease 35 | -0.373 | 7.0E-04 |
| ENSCGRG00001011886 | Sh2d3c | SH2 domain containing 3C | -0.364 | 5.6E-05 |
| ENSCGRG00001011912 | Retreg3 | reticulophagy regulator family member 3 | 0.361 | 7.8E-04 |
| ENSCGRG00001011944 | RCAN1 | Calcipressin-1 | -0.529 | 7.2E-18 |
| ENSCGRG00001011967 | Hexdc | hexosaminidase (glycosyl hydrolase family 20, catalytic domain) containing | 0.307 | 3.3E-06 |
| ENSCGRG00001012011 | Taf15 | TATA-box binding protein associated factor 15 | 0.399 | 6.3E-06 |
| ENSCGRG00001012092 | | histone H3 | 0.363 | 4.5E-06 |
| ENSCGRG00001012126 | Ercc6 | ERCC excision repair 6, chromatin remodeling factor | 0.310 | 5.4E-04 |
| ENSCGRG00001012408 | Glrx | glutaredoxin | -0.331 | 4.8E-11 |
| ENSCGRG00001012430 | Tbc1d5 | TBC1 domain family member 5 | 0.359 | 2.2E-07 |
| ENSCGRG00001012591 | Tmem260 | transmembrane protein 260 | 0.486 | 1.3E-06 |
| ENSCGRG00001012602 | Sema3e | semaphorin 3E | -0.333 | 5.9E-05 |
| ENSCGRG00001012679 | Kntc1 | kinetochore-associated protein 1 | 0.395 | 3.2E-08 |
| ENSCGRG00001012772 | Man1b1 | mannosidase, alpha, class 1B, member 1 | 0.332 | 1.0E-08 |
| ENSCGRG00001012914 | Ptpre | protein tyrosine phosphatase receptor type E | -0.642 | 4.1E-04 |
| ENSCGRG00001013038 | Ly6m | lymphocyte antigen 6D | -2.337 | 2.7E-05 |
| ENSCGRG00001013053 | Mmp12 | matrix metallopeptidase 12 | -1.122 | 7.7E-05 |
| ENSCGRG00001013097 | Hmox1 | heme oxygenase 1 | -0.376 | 7.7E-04 |
| ENSCGRG00001013303 | Tlr2 | toll-like receptor 2 | -0.695 | 7.5E-09 |
| ENSCGRG00001013389 | Eef1a2 | eukaryotic translation elongation factor 1 alpha 2 | 0.482 | 2.8E-05 |
| ENSCGRG00001013403 | | | 0.708 | 2.4E-04 |
| ENSCGRG00001013528 | Hdgf | heparin binding growth factor | -0.598 | 5.5E-38 |
| ENSCGRG00001013529 | Pibf1 | progesterone immunomodulatory binding factor 1 | 0.315 | 4.3E-04 |
| ENSCGRG00001013535 | Slc7a11 | solute carrier family 7 member 11 | -0.515 | 8.7E-04 |
| ENSCGRG00001013636 | Prss22 | brain-specific serine protease 4 | -0.851 | 9.9E-11 |
| ENSCGRG00001013659 | | histone H2B type 1 | 0.409 | 2.2E-05 |
| ENSCGRG00001013683 | | H-2 class I histocompatibility antigen, Q10 alpha chain | -0.556 | 2.4E-06 |
| ENSCGRG00001013733 | Col5a2 | collagen alpha-2(V) chain | -0.387 | 5.2E-05 |
| ENSCGRG00001013796 | Adgrg1 | adhesion G protein-coupled receptor G1 | -0.438 | 2.8E-14 |

| | | | | |
|---|---|---|---|---|
| ENSCGRG00001013909 | Gprc5a | G protein-coupled receptor class C group 5 member A | -0.382 | 4.0E-10 |
| ENSCGRG00001013914 | H6pd | hexose-6-phosphate dehydrogenase (glucose 1-dehydrogenase) | 0.313 | 1.2E-05 |
| ENSCGRG00001013957 | Wnt4 | Wnt family member 4 | -0.552 | 8.8E-08 |
| ENSCGRG00001013984 | Krt7 | keratin 7 | -0.618 | 7.4E-18 |
| ENSCGRG00001014022 | Poglut1 | protein O-glucosyltransferase 1 | 0.365 | 2.4E-08 |
| ENSCGRG00001014196 | Arhgap25 | Rho GTPase activating protein 25 | 0.365 | 3.6E-05 |
| ENSCGRG00001014213 | Cd82 | CD82 molecule | -1.154 | 3.0E-20 |
| ENSCGRG00001014220 | Rnf212 | ring finger protein 212 | 0.395 | 1.6E-04 |
| ENSCGRG00001014268 | Nemp1 | nuclear envelope integral membrane protein 1 | 0.427 | 2.5E-04 |
| ENSCGRG00001014386 | Nckap1l | NCK associated protein 1 like | -0.808 | 4.8E-04 |
| ENSCGRG00001014565 | St7l | suppression of tumorigenicity 7 like | 0.385 | 8.2E-04 |
| ENSCGRG00001014797 | | | -0.404 | 9.1E-08 |
| ENSCGRG00001014957 | Cit | citron rho-interacting serine/threonine kinase | 0.340 | 2.9E-07 |
| ENSCGRG00001014979 | Nupr1 | nuclear protein 1, transcriptional regulator | -0.463 | 9.1E-06 |
| ENSCGRG00001015046 | Rtel1 | regulator of telomere elongation helicase 1 | 0.320 | 1.5E-07 |
| ENSCGRG00001015135 | Junb | JunB proto-oncogene, AP-1 transcription factor subunit | -0.346 | 4.2E-05 |
| ENSCGRG00001015138 | Fam120c | family with sequence similarity 120C | 0.339 | 4.6E-05 |
| ENSCGRG00001015326 | Trrap | transformation/transcription domain-associated protein | 0.309 | 3.4E-04 |
| ENSCGRG00001015379 | Ubp1 | upstream binding protein 1 | 0.328 | 1.9E-06 |
| ENSCGRG00001015475 | Srpx2 | sushi repeat containing protein X-linked 2 | -0.560 | 5.6E-06 |
| ENSCGRG00001015519 | Mmp9 | matrix metallopeptidase 9 | -0.511 | 6.2E-08 |
| ENSCGRG00001015556 | Hspb1 | heat shock protein family B (small) member 1 | -0.346 | 3.0E-20 |
| ENSCGRG00001015625 | Atrnl1 | attractin like 1 | 0.313 | 2.5E-10 |
| ENSCGRG00001015703 | Txnrd1 | thioredoxin reductase 1 | -0.323 | 1.5E-04 |
| ENSCGRG00001015725 | Gch1 | GTP cyclohydrolase 1 | -0.322 | 1.4E-06 |
| ENSCGRG00001015794 | Gstm1 | glutathione S-transferase Y1 | -0.300 | 1.0E-18 |
| ENSCGRG00001015887 | Megf8 | multiple EGF-like-domains 8 | 0.356 | 6.5E-06 |
| ENSCGRG00001015969 | Fgfr2 | fibroblast growth factor receptor 2 | -0.400 | 8.1E-04 |
| ENSCGRG00001016081 | Metrnl | meteorin like, glial cell differentiation regulator | -0.361 | 9.0E-04 |
| ENSCGRG00001016112 | Gas7 | growth arrest specific 7 | -0.855 | 3.8E-10 |
| ENSCGRG00001016193 | Pank1 | pantothenate kinase 1 | -0.366 | 6.2E-04 |
| ENSCGRG00001016280 | Slpi | secretory leukocyte peptidase inhibitor | -0.541 | 9.2E-04 |
| ENSCGRG00001016315 | Tyk2 | tyrosine kinase 2 | 0.321 | 2.4E-06 |
| ENSCGRG00001016376 | Uggt2 | UDP-glucose glycoprotein glucosyltransferase 2 | 0.426 | 7.8E-06 |
| ENSCGRG00001016491 | Wrn | Werner syndrome RecQ like helicase | 0.334 | 8.9E-10 |
| ENSCGRG00001016581 | | placenta expressed transcript 1 | -0.481 | 7.4E-18 |
| ENSCGRG00001016633 | Syne2 | spectrin repeat containing nuclear envelope protein 2 | 0.363 | 8.0E-05 |

| | | | | |
|---|---|---|---|---|
| ENSCGRG00001016721 | S100a4 | S100 calcium binding protein A4 | -1.220 | 6.2E-27 |
| ENSCGRG00001016767 | Klhl4 | kelch like family member 4 | -0.370 | 3.4E-05 |
| ENSCGRG00001016824 | Arhgap33 | Rho GTPase activating protein 33 | 0.405 | 5.3E-06 |
| ENSCGRG00001016880 | Sfrp4 | secreted frizzled related protein 4 | -0.310 | 9.7E-04 |
| ENSCGRG00001016927 | S100a3 | S100 calcium binding protein A3 | -1.091 | 3.3E-06 |
| ENSCGRG00001017011 | Cdkl2 | cyclin dependent kinase like 2 | 0.399 | 1.6E-04 |
| ENSCGRG00001017065 | Plce1 | phospholipase C, epsilon 1 | 0.323 | 9.8E-04 |
| ENSCGRG00001017095 | Fads3 | fatty acid desaturase 3 | -0.346 | 4.6E-10 |
| ENSCGRG00001017137 | Pde1b | phosphodiesterase 1B | -0.492 | 7.9E-06 |
| ENSCGRG00001017362 | Rims2 | regulating synaptic membrane exocytosis 2 | 0.326 | 7.6E-04 |
| ENSCGRG00001017432 | Rpgr | sushi repeat containing protein X-linked | -0.346 | 1.8E-04 |
| ENSCGRG00001017589 | Ncoa5 | nuclear receptor coactivator 5 | 0.385 | 3.1E-09 |
| ENSCGRG00001017591 | | | -0.587 | 1.2E-06 |
| ENSCGRG00001017614 | Pcnt | pericentrin (kendrin) | 0.305 | 2.9E-05 |
| ENSCGRG00001017681 | Tsen2 | tRNA splicing endonuclease subunit 2 | 0.307 | 5.3E-04 |
| ENSCGRG00001017691 | Itga5 | integrin alpha 5 (fibronectin receptor alpha) | -0.391 | 3.8E-05 |
| ENSCGRG00001017714 | Aldh1a1 | aldehyde dehydrogenase family 1, subfamily A1 | -0.612 | 3.3E-04 |
| ENSCGRG00001017847 | Loxl1 | lysyl oxidase like 1 | -0.401 | 2.9E-07 |
| ENSCGRG00001017915 | Dcxr | dicarbonyl and L-xylulose reductase | -0.423 | 5.3E-13 |
| ENSCGRG00001018092 | | septin 3 | -0.392 | 1.2E-04 |
| ENSCGRG00001018111 | AKR1B8 | Aldose reductase-related protein 2 | -0.302 | 1.9E-05 |
| ENSCGRG00001018120 | Mical2 | microtubule associated monooxygenase, calponin and LIM domain containing 2 | -0.365 | 2.8E-13 |
| ENSCGRG00001018155 | Sparc | secreted protein acidic and cysteine rich | -0.399 | 3.2E-04 |
| ENSCGRG00001018176 | Pde2a | phosphodiesterase 2A | -0.751 | 4.8E-13 |
| ENSCGRG00001018280 | Eml6 | EMAP like 6 | 0.447 | 2.7E-07 |
| ENSCGRG00001018320 | Emc1 | ER membrane protein complex subunit 1 | 0.300 | 9.7E-16 |
| ENSCGRG00001018370 | Angpt4 | angiopoietin 4 | -0.691 | 1.3E-08 |
| ENSCGRG00001018373 | Mab21l3 | mab-21 like 3 | -0.594 | 9.3E-05 |
| ENSCGRG00001018405 | Ahcyl1 | adenosylhomocysteinase like 1 | -0.361 | 8.9E-21 |
| ENSCGRG00001018466 | Gstm7 | glutathione S-transferase Mu 7 | -0.398 | 1.3E-05 |
| ENSCGRG00001018587 | Plcxd2 | phosphatidylinositol specific phospholipase C X domain containing 2 | -0.459 | 6.2E-09 |
| ENSCGRG00001018610 | Ext2 | exostosin glycosyltransferase 2 | 0.305 | 4.5E-08 |
| ENSCGRG00001018659 | Vps41 | VPS41 subunit of HOPS complex | 0.306 | 5.5E-08 |
| ENSCGRG00001018660 | Mphosph9 | M-phase phosphoprotein 9 | 0.364 | 6.2E-07 |
| ENSCGRG00001018874 | | | -0.305 | 3.9E-04 |
| ENSCGRG00001018914 | Nqo1 | NAD(P)H quinone dehydrogenase 1 | -0.616 | 9.1E-05 |
| ENSCGRG00001018933 | Glg1 | golgi apparatus protein 1 | 0.327 | 2.3E-08 |
| ENSCGRG00001018973 | Zfand2a | zinc finger AN1-type containing 2A | -0.406 | 1.3E-04 |
| ENSCGRG00001019005 | Slc6a6 | solute carrier family 6 member 6 | -0.361 | 3.2E-04 |
| ENSCGRG00001019043 | Apbb3 | amyloid beta precursor protein binding family B member 3 | 0.480 | 9.0E-04 |

| | | | | |
|---|---|---|---|---|
| ENSCGRG00001019058 | Elmo1 | engulfment and cell motility 1 | -0.308 | 3.0E-05 |
| ENSCGRG00001019163 | | selenium binding protein 1 | -0.601 | 4.3E-06 |
| ENSCGRG00001019222 | Rhbdl2 | rhomboid like 2 | -0.681 | 3.3E-04 |
| ENSCGRG00001019357 | Ankrd1 | ankyrin repeat domain 1 (cardiac muscle) | -0.782 | 5.1E-13 |
| ENSCGRG00001019377 | Adgrd1 | adhesion G protein-coupled receptor D1 | -0.472 | 3.4E-16 |
| ENSCGRG00001019430 | Ldhd | lactate dehydrogenase D | -0.424 | 1.1E-04 |
| ENSCGRG00001019560 | Map1a | microtubule-associated protein 1 A | -0.426 | 1.5E-09 |
| ENSCGRG00001019565 | Nbeal2 | neurobeachin like 2 | 0.337 | 1.8E-05 |
| ENSCGRG00001019571 | Kif15 | kinesin family member 15 | 0.320 | 9.0E-05 |
| ENSCGRG00001019644 | Ccn5 | cellular communication network factor 5 | -0.895 | 2.7E-08 |
| ENSCGRG00001019687 | Abhd14a | abhydrolase domain containing 14A | 0.537 | 1.1E-04 |
| ENSCGRG00001019694 | Fhl2 | four and a half LIM domains 2 | -0.388 | 2.1E-08 |
| ENSCGRG00001019750 | Pdpn | podoplanin | -0.370 | 2.0E-05 |
| ENSCGRG00001019785 | Zfp362 | zinc finger protein 362 | 0.441 | 2.0E-08 |
| ENSCGRG00001019795 | Dcn | decorin | -0.441 | 6.4E-10 |
| ENSCGRG00001020100 | Oplah | 5-oxoprolinase, ATP-hydrolysing | 0.339 | 3.7E-05 |
| ENSCGRG00001020129 | Phka1 | phosphorylase kinase regulatory subunit alpha 1 | 0.342 | 6.5E-06 |
| ENSCGRG00001020144 | Anxa1 | annexin A1 | -0.326 | 3.5E-11 |
| ENSCGRG00001020266 | Man1a2 | mannosidase, alpha, class 1A, member 2 | 0.390 | 2.5E-09 |
| ENSCGRG00001020314 | Tmem108 | transmembrane protein 108 | -0.619 | 1.0E-06 |
| ENSCGRG00001020324 | Pkd1 | polycystin 1, transient receptor poteintial channel interacting | 0.440 | 9.6E-17 |
| ENSCGRG00001020328 | Cyth2 | cytohesin 2 | 0.414 | 2.3E-06 |
| ENSCGRG00001020376 | Ska3 | spindle and kinetochore associated complex subunit 3 | 0.339 | 1.1E-04 |
| ENSCGRG00001020442 | 1500009L16Rik | RIKEN cDNA 1500009L16 gene | -0.351 | 1.8E-07 |
| ENSCGRG00001020445 | | hydroxypyruvate isomerase (putative) | -0.304 | 2.2E-04 |
| ENSCGRG00001020497 | Pm20d2 | peptidase M20 domain containing 2 | 0.408 | 3.6E-05 |
| ENSCGRG00001020544 | Spdl1 | spindle apparatus coiled-coil protein 1 | 0.390 | 8.0E-04 |
| ENSCGRG00001020582 | Zfp523 | zinc finger protein 76 | 0.397 | 1.9E-07 |
| ENSCGRG00001020635 | | septin 4 | -0.461 | 7.4E-04 |
| ENSCGRG00001020680 | Scarf2 | scavenger receptor class F member 2 | 0.402 | 4.7E-06 |
| ENSCGRG00001020688 | Rogdi | rogdi atypical leucine zipper | -0.314 | 3.2E-04 |
| ENSCGRG00001020696 | Flot1 | flotillin 1 | -0.420 | 1.2E-10 |
| ENSCGRG00001020898 | Yae1d1 | YAE1 maturation factor of ABCE1 | -0.306 | 8.6E-05 |
| ENSCGRG00001020904 | Abcb8 | ATP binding cassette subfamily B member 8 | 0.350 | 7.0E-05 |
| ENSCGRG00001020961 | Atf5 | activating transcription factor 5 | 0.417 | 2.6E-06 |
| ENSCGRG00001020969 | DHDDS | dehydrodolichyl diphosphate synthase subunit | 0.630 | 7.6E-05 |
| ENSCGRG00001021007 | Hmgcs1 | 3-hydroxy-3-methylglutaryl-Coenzyme A synthase 1 | -0.353 | 2.4E-04 |
| ENSCGRG00001021074 | Axl | AXL receptor tyrosine kinase | -0.307 | 4.4E-06 |
| ENSCGRG00001021080 | Oasl1 | 2'-5' oligoadenylate synthetase-like 1 | -0.335 | 2.3E-05 |

| | | | | |
|---|---|---|---|---|
| ENSCGRG00001021083 | Iqgap3 | IQ motif containing GTPase activating protein 3 | 0.343 | 5.1E-09 |
| ENSCGRG00001021084 | Ltbp1 | latent transforming growth factor beta binding protein 1 | -0.402 | 4.1E-08 |
| ENSCGRG00001021122 | Rcan3 | RCAN family member 3 | 0.377 | 1.4E-06 |
| ENSCGRG00001021132 | Pla2g4a | phospholipase A2 group IVA | -0.363 | 4.0E-10 |
| ENSCGRG00001021180 | Loxl4 | lysyl oxidase like 4 | -0.576 | 2.4E-04 |
| ENSCGRG00001021189 | | von Willebrand factor A domain-containing protein 5A | -0.679 | 1.5E-09 |
| ENSCGRG00001021238 | Mrgprf | MAS related GPR family member F | -0.938 | 6.4E-04 |
| ENSCGRG00001021368 | Cpne2 | copine 2 | 0.308 | 5.0E-09 |
| ENSCGRG00001021379 | Acbd4 | acyl-CoA binding domain containing 4 | -0.408 | 1.5E-05 |
| ENSCGRG00001021452 | Gmppa | GDP-mannose pyrophosphorylase A | -0.318 | 7.3E-04 |
| ENSCGRG00001021524 | | ADAM metallopeptidase domain 19 | -0.308 | 2.2E-04 |
| ENSCGRG00001021545 | 6430571L13Rik | chromosome unknown C3orf18 homolog | -0.508 | 5.2E-10 |
| ENSCGRG00001021644 | Dyrk3 | dual specificity tyrosine phosphorylation regulated kinase 3 | -0.599 | 2.5E-11 |
| ENSCGRG00001021689 | Cnn1 | calponin 1 | -0.973 | 1.9E-05 |
| ENSCGRG00001021745 | Rfx3 | regulatory factor X3 | 0.371 | 4.5E-04 |
| ENSCGRG00001021785 | Prdx5 | peroxiredoxin 5 | -0.300 | 1.7E-05 |
| ENSCGRG00001021805 | Cpeb4 | cytoplasmic polyadenylation element binding protein 4 | -0.302 | 3.3E-04 |
| ENSCGRG00001021809 | Rusf1 | cDNA sequence BC017158 | -0.395 | 7.8E-09 |
| ENSCGRG00001021913 | Mgarp | mitochondria localized glutamic acid rich protein | -0.302 | 1.0E-05 |
| ENSCGRG00001022103 | SAT1 | Diamine acetyltransferase 1 | -0.337 | 8.9E-11 |
| ENSCGRG00001022116 | Galm | galactose mutarotase | -0.341 | 5.3E-07 |
| ENSCGRG00001022122 | Cited1 | Cbp/p300 interacting transactivator with Glu/Asp rich carboxy-terminal domain 1 | -0.426 | 2.9E-05 |
| ENSCGRG00001022275 | Dnmt1 | DNA methyltransferase (cytosine-5) 1 | 0.371 | 1.1E-11 |
| ENSCGRG00001022424 | Aldh4a1 | aldehyde dehydrogenase 4 family, member A1 | -0.350 | 3.3E-09 |
| ENSCGRG00001022444 | Ezr | ezrin | -0.343 | 6.5E-09 |
| ENSCGRG00001022515 | Fbp2 | fructose-bisphosphatase 2 | -0.441 | 8.6E-04 |
| ENSCGRG00001022612 | Arhgef25 | Rho guanine nucleotide exchange factor 25 | -0.918 | 4.0E-09 |
| ENSCGRG00001022792 | Afg2b | spermatogenesis associated 5-like 1 | -0.309 | 6.9E-06 |
| ENSCGRG00001022828 | Zfp113 | zinc finger protein 113 | 0.383 | 7.0E-04 |
| ENSCGRG00001022899 | Pir | pirin | -0.330 | 2.8E-04 |
| ENSCGRG00001022924 | Zc3h7b | zinc finger CCCH-type containing 7B | 0.319 | 8.4E-14 |
| ENSCGRG00001022969 | Iyd | iodotyrosine deiodinase | -0.497 | 8.0E-11 |
| ENSCGRG00001023035 | Mlph | melanophilin | 0.396 | 2.6E-04 |
| ENSCGRG00001023036 | Tiam2 | TIAM Rac1 associated GEF 2 | -0.323 | 9.2E-04 |
| ENSCGRG00001023083 | Limk1 | LIM domain kinase 1 | -0.324 | 1.1E-10 |
| ENSCGRG00001023123 | | testin LIM domain protein | -0.488 | 1.3E-11 |
| ENSCGRG00001023193 | Ltbp3 | latent transforming growth factor beta binding protein 3 | 0.507 | 4.6E-10 |

| | | | | |
|---|---|---|---|---|
| ENSCGRG00001023241 | Gss | glutathione synthetase | -0.347 | 5.0E-09 |
| ENSCGRG00001023280 | Plau | plasminogen activator, urokinase | 0.499 | 1.9E-04 |
| ENSCGRG00001023297 | Gmppb | GDP-mannose pyrophosphorylase B | -0.357 | 5.6E-06 |
| ENSCGRG00001023333 | Mamdc2 | MAM domain containing 2 | -0.379 | 1.3E-06 |
| ENSCGRG00001023432 | Cep128 | centrosomal protein 128 | 0.338 | 9.3E-05 |
| ENSCGRG00001023457 | Thbs2 | thrombospondin 2 | -0.427 | 3.4E-08 |
| ENSCGRG00001023716 | Fchsd1 | FCH and double SH3 domains 1 | -0.324 | 1.0E-04 |
| ENSCGRG00001023800 | Pycr1 | pyrroline-5-carboxylate reductase 1 | -0.418 | 4.5E-05 |
| ENSCGRG00001023801 | Cda | cytidine deaminase | -0.448 | 2.3E-06 |
| ENSCGRG00001023903 | Haus8 | HAUS augmin like complex subunit 8 | 0.346 | 8.5E-04 |
| ENSCGRG00001023905 | Btbd3 | BTB domain containing 3 | -0.336 | 2.4E-06 |
| ENSCGRG00001024012 | Itga7 | integrin subunit alpha 7 | -0.535 | 1.5E-05 |
| ENSCGRG00001024045 | Cnksr1 | connector enhancer of kinase suppressor of Ras 1 | -0.388 | 8.8E-04 |
| ENSCGRG00001024096 | Slc25a1 | solute carrier family 25 member 1 | -0.348 | 5.4E-04 |
| ENSCGRG00001024132 | | Rho guanine nucleotide exchange factor 4 | -0.521 | 7.6E-06 |
| ENSCGRG00001024135 | Il6 | interleukin 6 | -0.555 | 4.3E-04 |
| ENSCGRG00001024156 | Nagpa | N-acetylglucosamine-1-phosphodiester alpha-N-acetylglucosaminidase | -0.300 | 1.1E-04 |
| ENSCGRG00001024160 | Fblim1 | filamin binding LIM protein 1 | -0.366 | 1.7E-04 |
| ENSCGRG00001024241 | | microtubule associated scaffold protein 1 | -0.440 | 5.4E-06 |
| ENSCGRG00001024332 | Limd2 | LIM domain containing 2 | -0.365 | 3.5E-04 |
| ENSCGRG00001024401 | Plpp1 | phospholipid phosphatase 1 | -0.350 | 2.3E-04 |
| ENSCGRG00001024553 | Fam161b | family with sequence similarity 161, member B | -0.355 | 8.4E-04 |
| ENSCGRG00001024694 | Peak1 | pseudopodium-enriched atypical kinase 1 | 0.359 | 2.4E-04 |
| ENSCGRG00001024710 | Scly | selenocysteine lyase | -0.331 | 1.1E-05 |
| ENSCGRG00001024723 | Asb1 | ankyrin repeat and SOCS box containing 1 | -0.417 | 8.1E-04 |
| ENSCGRG00001024752 | Tomm40l | translocase of outer mitochondrial membrane 40 like | -0.306 | 1.4E-06 |
| ENSCGRG00001024863 | Ehd4 | EH domain containing 4 | -0.330 | 6.9E-10 |
| ENSCGRG00001024963 | Mpv17 | MpV17 mitochondrial inner membrane protein | -0.315 | 7.5E-04 |
| ENSCGRG00001024973 | Scap | Cricetulus griseus SREBF chaperone (Scap), mRNA. | 0.332 | 1.2E-05 |
| ENSCGRG00001025003 | Fdps | farnesyl diphosphate synthase | -0.357 | 1.0E-04 |
| ENSCGRG00001025027 | Fcsk | fucose kinase | 0.343 | 4.1E-06 |

#### Table S2: Gene set enrichment analysis results

*Table S2: GSEA analysis for the differentially expressed genes between control and NAM-fed conditions. The reference gene set was WikiPathways Human 2024.*

| Term | ES | NES | NOM p-val | FDR q-val | FWER p-val | Lead_genes |
|---|---|---|---|---|---|---|
| Oxidative Phosphorylation WP623 | 0.88 | 2.64 | 0 | 0 | 0 | ND1, ND2, ND3, ND5, ND4L, ND4, ATP6 |
| Electron Transport Chain OXPHOS System In Mitochondria WP111 | 0.89 | 2.88 | 0 | 0 | 0 | ND1, ND2, ND3, ND5, CYTB, ND4L, ND4 |
| Mitochondrial Complex I Assembly Model OXPHOS System WP4324 | 0.97 | 2.50 | 0 | 0 | 0 | ND1, ND2, ND5, ND4L, ND4 |
| PI3K Akt Signaling WP4172 | -0.61 | -1.96 | 0.0044 | 0.0491 | 0.034 | SGK2, TLR2, ANGPT4, IL6, LAMC2, ITGA7, CREB3L1, THBS2, FGFR2, ITGA5 |

#### Figure S4: DESeq2 comparison between 2mM and 5mM NAM feeds

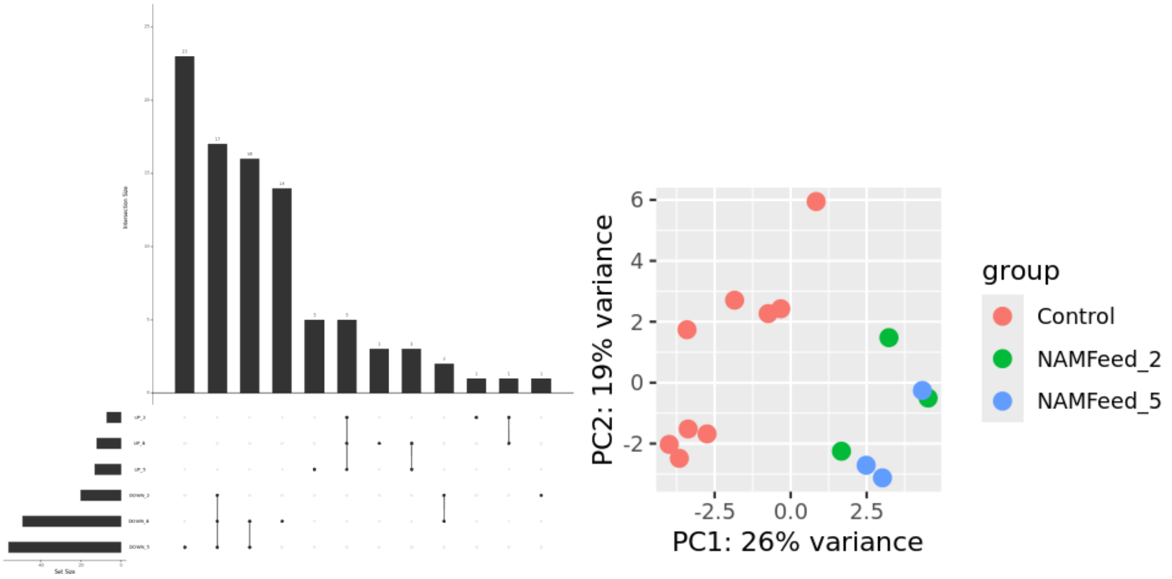


*Figure S4. Gene expression comparison between 2mM and 5mM NAM feeds. A) UpSet plot showing intersections of differentially expressed genes across NAM feed conditions. Sets represent upregulated (UP) and downregulated (DOWN) genes at 2mM, 5mM, and both (B) NAM concentrations. The bar plot above shows the size of each gene intersection, while the set size bars on the left indicate total genes per condition. B) PCA plot showing the distribution of log2 (TPM+1) gene expression comparing control versus 2mM and 5mM NAM feeds.*
